# DCHS1 Modulates Forebrain Proportions in Modern Humans via a Glycosylation Change

**DOI:** 10.1101/2025.05.14.654031

**Authors:** M. Veronica Pravata, Andrea Forero, Ane C. Ayo Martin, Giovanna Berto, Tim Heymann, Luise Fast, Matthias Mann, Stephan Riesenberg, Silvia Cappello

## Abstract

Comparative anatomical studies of primates and extinct hominins, including Neanderthals, show that the modern human brain is characterised by a disproportionately enlarged neocortex relative to the striatum. To explore the molecular basis of this difference, we screened for missense mutations that are unique to modern humans and occur at high frequency and that alter post-translational sites. One such mutation was identified in *DCHS1*, a protocadherin family gene, and it was found to disrupt an N-glycosylation site in modern humans. Using CRISPR/Cas9-editing we introduced into human-induced pluripotent stem cells (hiPSCs) this ancestral *DCHS1* variant present in Neanderthals and other primates, representing the ancestral state before the modern human-specific substitution. Leveraging hiPSCs-derived neural organoids, we observed an expansion of striatal progenitors at the expense of the neocortex, mirroring the anatomical distribution seen in non-human primates. We further identify the ephrin receptor EPHA4 as a binding partner of DCHS1 and show that modern human-specific alterations in DCHS1 modulate EPHA4-ephrin signalling, contributing to a gradual shift in the neocortex-to-striatum ratio - a hallmark of brain organisation in our species.

## Main

Sociality and cognition represent areas in which modern humans may differ significantly from other hominins and primates. Comparisons with the great apes, our closest living relatives, have provided crucial insights into the evolution of human behavior and cognition^1-5^. Anatomical evidence, including volumetric measurements and endocast analyses, suggests that the reorganization and expansion of the neocortex played a pivotal role in shaping our species’ advanced cognitive and social capacities ^4,6,7^.

Recent findings indicate that modern human brain evolution involved a redistribution of neural resources rather than simple brain enlargement ^8,9^. Despite similar overall brain volumes, Homo sapiens exhibited an expanded cerebral cortex at the expense of subcortical structures, a shift that distinguishes us from Neanderthals. This cortex-striatum trade-off, evident across primate evolution, may have enhanced cognitive flexibility and social networking in modern humans, potentially at the cost of the robust habit formation and procedural memory systems that characterize other primates and earlier hominins.

Understanding the genetic basis of these anatomical shifts is now within reach, thanks to comparative genomic analyses of late hominins. The availability of high-quality genomes from Neanderthals and Denisovans allows us to identify genetic changes unique to modern humans, providing a molecular framework to investigate brain reorganization. Among these, modern human–specific single nucleotide variants (SNVs) have emerged as promising candidates for shaping these developmental differences. The identification of novel genetic changes shared by nearly all modern humans can shed light on the molecular events that fine-tuned modern human physiology, including brain development. Comparative studies with archaic humans have revealed several genetic variants in the modern human lineage, such as insertions, deletions, and SNVs ^10-15^. Notably, recent analyses have shown that genes harboring modern human-specific mutations show enriched expression in the orbital frontal cortex specifically during infancy (0-2 years), whereas genes harboring archaic human-specific mutations show no such enrichment^14^. Additionally, mutations in genes such as *KIF18A, KNL1, TKTL1, CHD2*, and *NOVA1* have been shown to influence neuronal progenitor populations during early brain development^16-22^, collectively suggesting cellular mechanisms that may have contributed to cortical expansion.

Here, we investigate the disproportionate expansion of the neocortex relative to the striatum, a defining anatomical feature that distinguishes modern humans from our ancestors. In search of the genetic basis of this shift, we screened for modern human-specific missense mutations absent in archaic humans and identified a SNV in *Dachsous Cadherin-Related 1 (DCHS1)*, which disrupts a conserved N-glycosylation site. Using CRISPR/Cas9 to introduce the ancestral allele into hiPSCs and comparative analyses of iPSC-derived neural organoids, we observed a shift in the balance of neocortical and striatal progenitors in organoids harboring ancestral *DCHS1*. Single-cell RNA sequencing (scRNA-seq) and biochemical assays revealed that these differences may be driven by altered interactions between DCHS1 and its newly identified binding partner ephrin type-A receptor 4 (EPHA4). This work provides evidence that a post-translational modification driven by a modern human-specific SNV can influence brain development, potentially contributing to distinctive anatomical features of our species.

## Results

### Modern humans have an increased neocortex-to-striatum ratio compared to other primates

The overall shape of the primate brain has changed considerably during evolution, in part due to increasing encephalization. Several lines of evidence suggest that neocortical expansion and reorganization began before the emergence of hominins, potentially laying the groundwork for more pronounced changes in later species. To investigate how these shifts manifest in modern humans relative to other primates, we compiled previously published volumetric data and conducted additional measurements^23^. We quantified relative volumes of various brain regions and calculated percentage differences between modern humans and other primates (Fig. 1a-d) (Supplementary Figs. 1c).

**Fig. 1.**
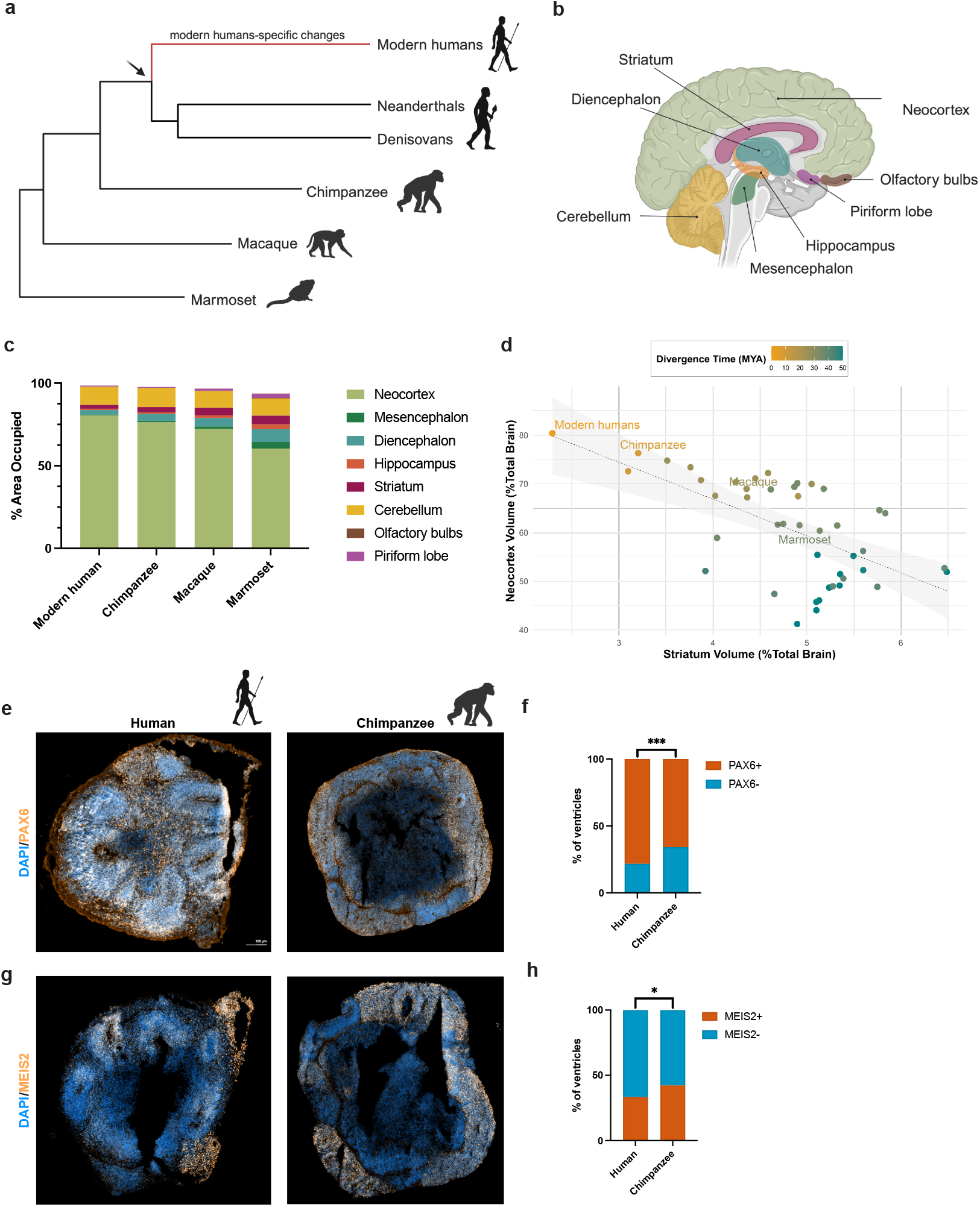
| Modern humans have an increased neocortex-to-striatum ratio compared to other primates. (a) Schematic representation of evolutionary relationships among modern humans and closely related species, including Neanderthals, Denisovans, chimpanzees, macaques, and marmosets. Created in https://BioRender.com. **(b)** Illustration of the human brain, highlighting key brain regions analyzed for volumetric comparisons across species. Created in https://BioRender.com. **(c)** Stacked bar plots showing the relative volumetric proportions of the neocortex, mesencephalon, diencephalon, hippocampus, striatum, cerebellum, olfactory bulbs, and piriform lobe in present-day humans, chimpanzees, macaques, and marmosets. Data were extrapolated from Frahm et al. (1981). **(d)** Relationship between striatum and neocortex volumes across primate species. A linear regression was performed to examine the relationship between striatum volume (% of total brain volume) (X-axis) and neocortex volume (% of total brain volume) (Y-axis) across primates. Each point represents a species, color-coded by divergence time (million years ago, MYA), with earlier-diverging species shown in orange and more recent divergences in teal. Data were extrapolated from Frahm et al. (1981). Representative immunohistochemistry (IHC) images of PAX6+ **(e)** and MEIS2+ **(g)** cells in 30-day-old neural organoids derived from human and chimpanzee iPSCs, along with quantification. Statistical significance was assessed using a binomial test by comparing the chimpanzee condition with each human condition individually. **(f)** For PAX6 (Batch = 1), n = 4 organoids per condition; total number of ventricles: Human = 23, Chimpanzee = 73. **(f)** For MEIS2 (Batch = 1), n = 4 organoids per condition; total number of ventricles: Human = 30, Chimpanzee = 52.

Humans had a larger neocortex than all other species, with the smallest difference compared to chimpanzees (4%), followed by macaques (8.2%), and the largest difference compared to marmosets (20%). Concomitantly, humans showed a smaller striatum, registering reductions of 0.9%, 2.3% and 2.8% relative to chimpanzees, macaques and marmosets, respectively. Although other brain structures in humans were also marginally smaller, these differences were modest compared to the change in the striatum (Supplementary Figs. 1c). Also, analysis of endocranial shapes showed that parts of the prefrontal cortex are relatively larger in modern humans than in Neanderthals ^9^. These findings suggest that the modern human neocortex expanded at the expense of subcortical regions, potentially laying the foundation for unique cognitive adaptations.

To validate these anatomical observations, we generated neural organoids from modern human and chimpanzee iPSCs and examined the distribution of cortical (PAX6+) versus striatal (MEIS2+) progenitor regions. In accordance with the volumetric data, human organoids showed a greater proportion of PAX6+ areas (12.5% difference), accompanied by fewer MEIS2+ regions (9% difference) compared to chimpanzee organoids (Fig. 1e-h). These results further support a human-specific shift in neocortex-to-striatum proportions, which may underlie distinctive cognitive and behavioral traits of our species.

### Modern human–specific missense SNVs can cause alterations in post-translational modifications

Based on the anatomical findings and observed differences in human and chimpanzee organoids, we hypothesized that the shift in neocortex-to-striatum proportions may stem from genetic mutations that arose after the split from the last common ancestor of modern humans and Neanderthals. To find potentially responsible genes, we first identified 83 genes carrying frequent missense variants unique to present-day humans (see https://bioinf.eva.mpg.de/catalogbrowser/protein-changing-variants) and evaluated their expression in the developing human brain using scRNA-seq data obtained from 5 to 14 post-conceptional weeks (PCW) ^24^. We mapped each gene’s expression across multiple cell types—radial glia, neural crest, intermediate progenitors (IPCs), immune cells, neurons, and neuroblasts—in both the dorsal forebrain (Excitatory) and the ganglionic eminences (GEs), which give rise to most inhibitory neurons (Inhibitory) and lateral GE– derived (LGE) striatal cells (Striatum) (Fig. 2a). For each gene, we calculated its mean expression level and percentile rank among all genes. Of the 83 candidates, 56 surpassed the 50th-percentile mark, while the remainder exhibited lower to negligible expression (Fig. 2a, Supplementary Figs. 1a). We also examined how these genes were distributed among specific clusters (Supplementary Figs. 1a,b). Some, such as *DCHS1* and *NOVA1*, were widely expressed across multiple cell types, whereas others exhibited more restricted patterns. These findings align with previous observations that certain genes (*TKTL1, KIF18A*) display notably high expression in specific cell types (e.g., neural crest, radial glia, neuronal IPCs). Of note, a few genes (e.g., *AHR* and *ZNF185*) were expressed predominantly in the dorsal compartment (Supplementary Figs. 1b), resulting in a specific regional pattern of expression.

**Fig. 2.**
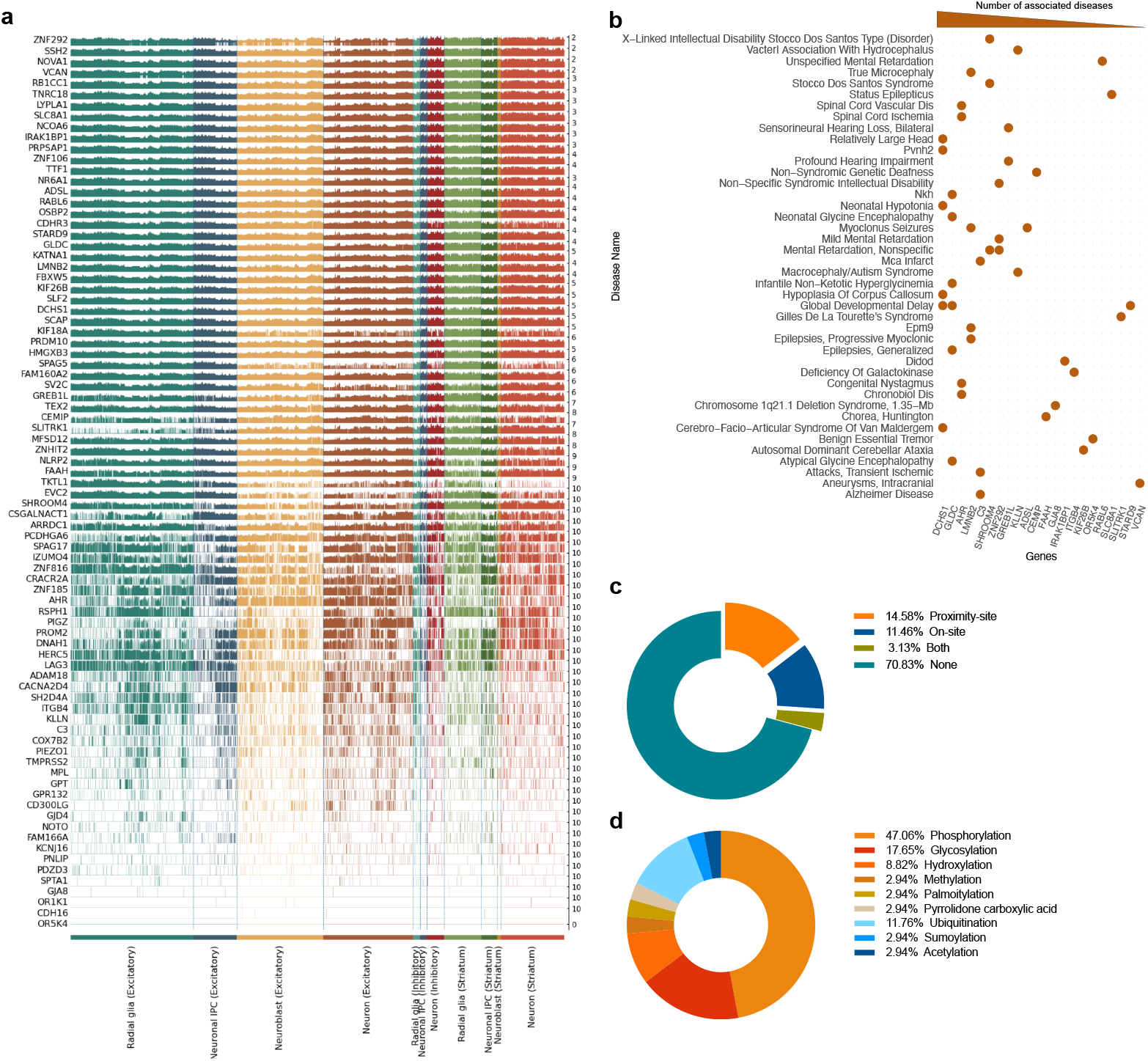
Unique modern human missense variants alter the post-translational modification (PTM) landscape. **a)** Expression levels of 83 genes associated with non-synonymous variants in modern humans. Data were obtained from single-cell RNA sequencing (scRNA-seq) of first-trimester human brain cells, including radial glia, neuronal intermediate progenitor cells (IPCs), neuroblasts, and excitatory, inhibitory, and striatal neurons. **(b)** Heatmap of genes linked to neurodevelopmental disorders, based on data from the DisGeNET database, highlighting their associations with specific disorders. **(c)** Pie chart depicting the distribution of mutations affecting PTM sites, categorized as *On-site* (directly at the modification site), *Proximity-site* (near the modification site), *Both* (affecting both the modification site and neighboring sites), or *None* (no effect on modification sites). **(d)** Breakdown of PTM types present at sites where modifications were identified, illustrating the diversity of PTMs affected by the variants.

In the next step, we mined DisGeNET ^25,26^ to determine whether these genes were associated with neurological disorders, reasoning that variants implicated in disease may also influence brain development. Notably, 22 genes were connected to various neurological conditions (Fig. 2b), including *ADSL, AHR*, and *SLITRK1*, for which modern-human specific phenotypes have been described previously^27-29^. *GLDC* and *DCHS1* were associated with the most neurodevelopmental disorders, suggesting crucial functions in brain development. In particular, *DCHS1* was also connected to craniofacial malformations, further hinting at possible effects on morphological traits. Given that there are significant craniofacial differences between modern humans and Neanderthals, with developmental changes occurring both pre- and postnatally^30-32^, genes such as *DCHS1* may have played a role in shaping species-specific facial morphology.

To determine whether these genes exhibit interspecies differences in expression, we turned to existing brain organoid datasets derived from macaque, chimpanzee, and hiPSCs. However, no marked differences were detected, indicating that transcriptional level changes alone may not fully account for modern human–specific neurodevelopmental patterns (Supplementary Figs. 1d-f).

We therefore set out to investigate post-translational modifications (PTMs), which play a crucial role in shaping protein conformation, stability, and interactions - key factors in developmental signaling networks^33-35^. To explore putative functional changes mediated by PTMs, we employed in silico predictions to identify major PTM types and assessed whether missense variants in modern humans could alter PTMs at the mutation site (“on-site”), in adjacent residues (“proximity”), or both (“both”) (Fig. 2c)^36-38^. Notably, more than a quarter of the missense mutations analyzed were predicted to induce PTM changes (Fig. 2c). Of these, half led to the putative loss or addition of a PTM at the site of mutation, while the other half resulted in putative PTM changes at neighboring amino acids, or both. Phosphorylation and glycosylation were the most common forms of PTMs involved (Fig. 2d), in line with that these are among the most common PTMs^39^. Collectively, these results suggest that changes in post-translational regulation could underlie some of the neurodevelopmental differences that set modern humans apart from earlier hominids and other primates.

### The modern human DCHS1 has lost a glycosylation site

Among the genes analyzed, *DCHS1* emerged as a particularly compelling candidate for several reasons. First, it was one of the few genes that showed strong and widespread expression in multiple cell types in the developing human brain, including radial glia, neural crest, and neuronal progenitors-key populations involved in cortical expansion and neurodevelopmental patterning. Second, *DCHS1* was associated with the most neurological disorders in DisGeNET, including both neurodevelopmental disorders and craniofacial malformations ^40-43^, suggesting a dual role in shaping brain and facial morphology. Given that modern humans exhibit an expanded neocortex-to-striatum ratio and distinct craniofacial morphology relative to Neanderthals, we hypothesized that species-specific changes in DCHS1 may have contributed to these evolutionary differences. Third, our PTM prediction identified a missense mutation in modern humans that disrupts an N-glycosylation site in DCHS1, a modification that can influence protein folding, stability, and receptor interactions. Together, these reasons position DCHS1 as a promising candidate for further investigation, particularly with respect to its role in developmental brain evolution.

*DCHS1* encodes a calcium-dependent adhesion molecule of the protocadherin superfamily that is linked to the Hippo signaling pathway, key for organ size, and planar cell polarity, well known for regulating morphology as well as neural progenitor proliferation and migration in mammals^40,41,43-45^. The non-synonymous substitution from an ancestral asparagine (Asn) to an aspartic acid (Asp) in modern humans affects DCHS1 in position 777, which carries an Asn residue in all primates sequenced to date (Fig. 3a). Comparative analysis across vertebrates, including primates, rodents, and other mammals, further supports conservation at this position (Supplementary Figs. 2a). Moreover, in silico prediction of the structural impact of the Asp at position 777 does not indicate structural effects on the overall protein structure (Prediction Score 0.000, v2.2.3r406) ^46^.

**Fig. 3.**
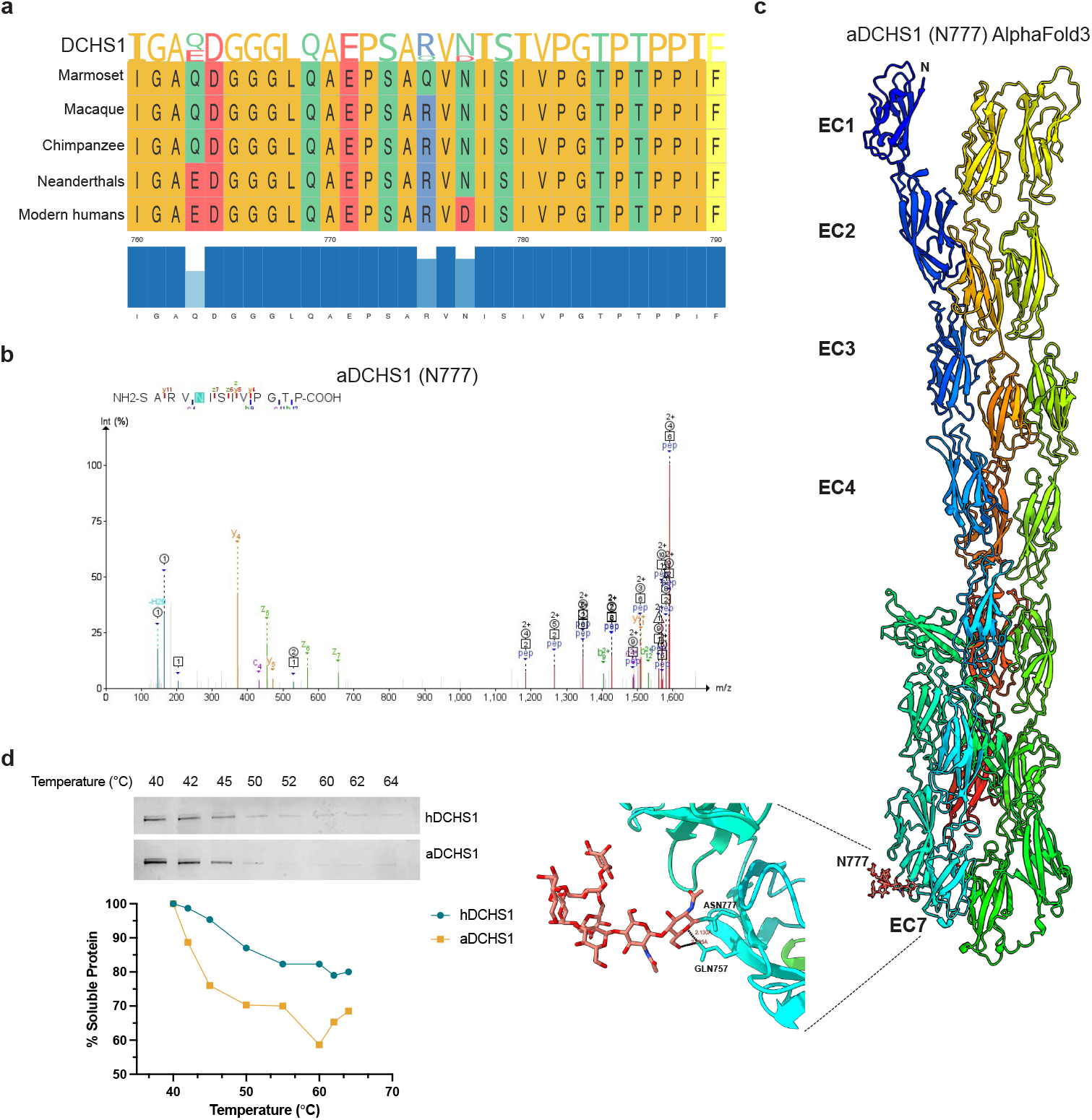
The modern DCHS1 variant leads to loss of glycosylation at position 777. **(a)** Sequence alignment of the region encompassing the D777N mutation across primates. **(b)** Mass spectrometry (MS) spectra showing glycosylation of aDCHS1 in the peptide *SARVNISIVPGTP*. **(c)** AlphaFold3-predicted structure of the aDCHS1 extracellular protocadherin (EC) domain, including glycosylation at position 777 and zoomed-in view highlighting the putative N-linked oligomannose at asparagine 777 in aDCHS1. **(d)** Quantification of CTSA assay based on western blot analysis of DCHS1 (n = 3). Solid lines represent the best fit of the data to a Boltzmann sigmoid curve in GraphPad Prism.

Asn777 resides within the Asn-X-Ser motif for N-linked glycosylation, a common post-translational modification. In silico analysis confirmed that while the ancestral DCHS1 (aDCHS1), present in Neanderthals and other primates, could undergo glycosylation at this site, the Asp777 substitution in modern humans disrupts this modification. Notably, all other predicted glycosylation sites remain unaffected by this change (Supplementary Figs. 2b,c). In light of this, we investigated the putative glycosylation by overexpressing the two forms of DCHS1 in human cells and analyzed their glycosylation by mass spectrometry (MS). This revealed that asparagine 777 in aDCHS1 carries an oligomannose glycan (Fig. 3b), while hDCHS1 does not (Supplementary Figs. 2d). We modeled the first 25 Cadherins of aDCHS1 using AlphaFold3 to inspect how Asp777 glycosylation could affect the overall structure (Fig. 3c). The ancestral glycosylation site is situated in the extracellular domain of DCHS1, specifically in the cadherin 7 (EC7) region, in close proximity to Gln757 and at the interface with the extracellular domain of cadherin 8 (Fig. 3c). Given the inherent conformational flexibility of protocadherins at each individual interface, this additional glycosylation site may serve to refine the overall binding affinity and stability of DCHS1^47-49^.

To investigate whether glycosylation might have an effect on the protein, we first investigated if this amino acid change could influence the thermal stability of DCHS1. We expressed the ancestral and modern human versions of the protein in HEK293 cells, which were then subjected to heating from 40°C to 64°C before cooling down, lysis and removal of denatured proteins by centrifugation^50^. DCHS1 protein that was not denatured was detected in the supernatant by western blotting. The results show that hDCHS1 exhibits greater thermal stability than aDCHS1, suggesting that the additional glycan branch might affect the flexibility of the protein (Fig. 3d).

### DCHS1 ancestralisation leads to increased striatum proportions

To investigate if the change in amino acid and glycosylation at position 777 results in altered neural phenotypes, we first used CRISPR genome editing to introduce the ancestral Asn residue into DCHS1 in hiPSCs, and then performed comparative analysis of iPSC-derived neural organoids carrying the ancestral or modern human version of DCHS1. To exclude potential CRISPR gRNA-dependent off-target effects that ultimately could result in phenotypic artefacts, we used two independent high-fidelity approaches using different gRNAs, high-fidelity Cas9 and Cas9D10A double nicking, to generate precisely edited clones (Supplementary Figs. 3a). As both approaches used different gRNAs, shared phenotypes observed from ancestralised cells cannot be due to potential gRNA-dependent off-target effects. We performed scRNA-seq of neural organoids at 30 and 60 days, when they are enriched for progenitors and neurons, respectively (Fig. 4a,c, Supplementary Figs. 3b,d). Uniform Manifold Approximation and Projection (UMAP) dimensionality reduction identified eight different cellular clusters (Fig. 4a,c), including neural progenitors, cortical neurons, and striatal projection neurons. To assess genotype-specific differences in neural development and differentiation, we examined the distribution of cell types in aDCHS1 and hDCHS1 at both 30 and 60 days (Fig. 4b,d) and calculated Cohen’s d effect size for each cell type (Supplementary Figs. 3c,e). The analysis revealed substantial differences in the frequencies of several cell types between the two genotypes. As expected, at 30 days most of the cells clustered in neural progenitor clusters, including cycling radial glia (CyRG), radial glia (RG), and dorsal radial glia (dRG) (Fig. 4b). Notably, aDCHS1 organoids exhibited a lower frequency of dRG (Cohen’s *d* Effect Size, d= 2.35) but a higher proportion of RG (d= -6.01), suggesting a redistribution within the progenitor population (Fig. 4b, Supplementary Figs. 3c). Pseudobulk RNA-seq analyses of the RG cluster revealed that hDCHS1 was enriched by dorsal forebrain markers, such as *TBR1, EMX2* and *HOPX*, while aDCHS1 RG mainly expressed more subpallial genes such as *RBFOX1, TAC1* and the striatal LGE marker *EBF1* (Supplementary Figs. 3f), suggesting that aDCHS1 progenitor possesses a more remarked subpallial profile compared to hDCHS1 RG.

**Fig. 4.**
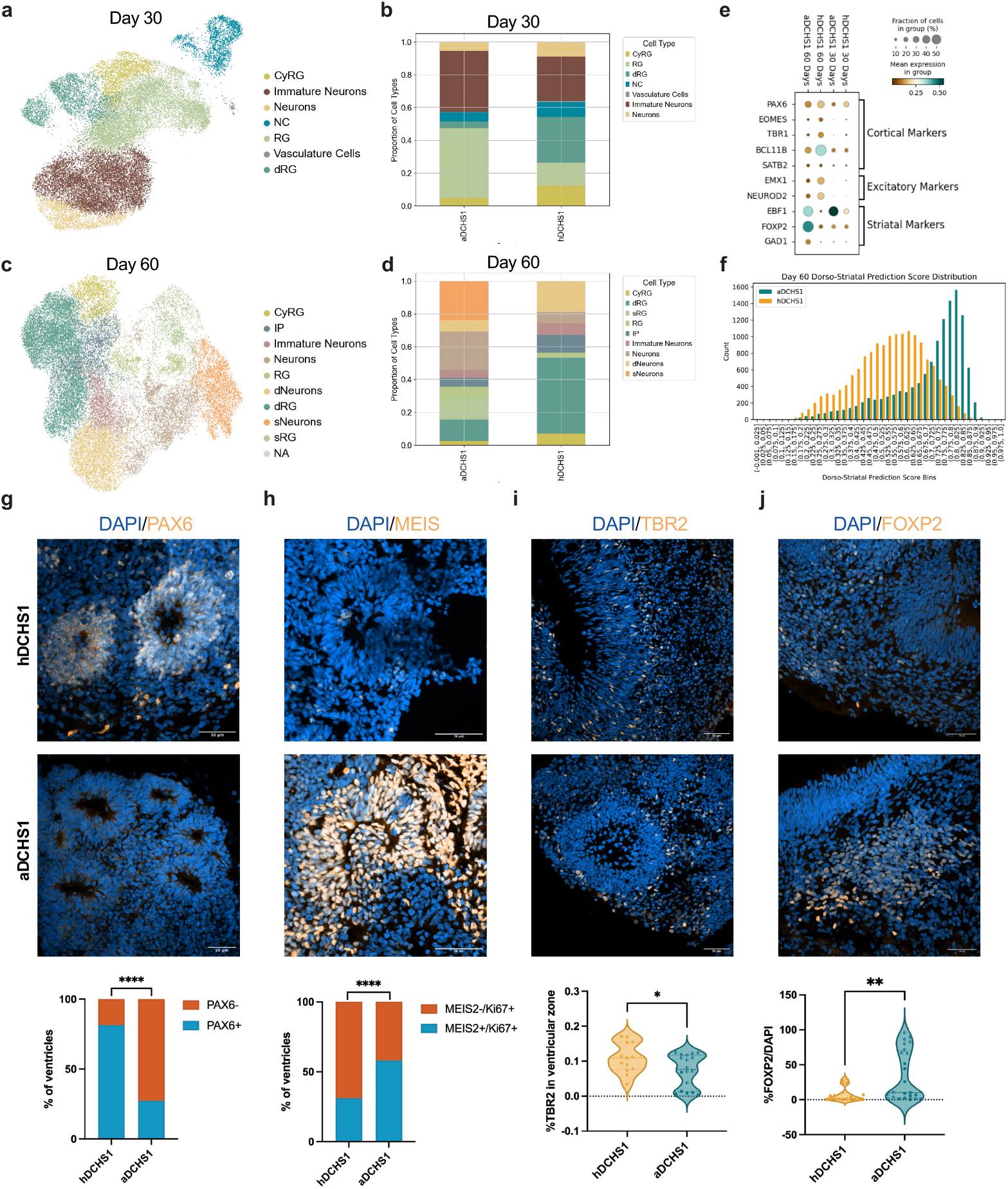
Neural organoids expressing ancestral and modern human DCHS1 exhibit different proportions of cortical and striatal lineages. **(a)** Uniform manifold approximation and projection (UMAP) visualization of single-cell RNA sequencing (scRNA-seq) clusters at day 30 in aDCHS1 and hDCHS1 neural organoids. Each color represents a distinct cell population. **(b)** Frequency analysis of scRNA-seq cell populations in day 30 aDCHS1 and hDCHS1 organoids. **(c)** UMAP visualization of scRNA-seq clusters at day 60 in aDCHS1 and hDCHS1 organoids. **(d)** Frequency analysis of scRNA-seq cell populations in day 60 aDCHS1 and hDCHS1 organoids. **(e)** Dot plot showing the expression of cortical, excitatory, and striatal marker genes. Dot color represents the average expression level, and dot size indicates the proportion of cells expressing each gene. **(f)** Histogram depicting the dorso-striatal (DV) score in aDCHS1 and hDCHS1 neural organoids at day 60. **(g-l)** Representative immunohistochemistry (IHC) images and quantification of lineage-specific markers: PAX6+ **(g)** and MEIS2+ **(h)** cells at day 30, and TBR2+ **(i)** and FOXP2+ **(j)** cells at day 60 in aDCHS1 and hDCHS1 neural organoids. Quantification data are shown as violin plots with the median represented by a dotted line. Each shape represents an individual cell line (n = number of analyzed ventricles from two independent batches of neural organoids). Statistical significance was assessed using one-way ANOVA. Scale bar: 50 µm.

At 60 days, the differences were more pronounced. hDCHS1 neural organoids showed higher frequencies of Intermediate Progenitors (Cohen’s *d* Effect Size, d= 0.95) and dorsal Neurons (dNeurons, d=1.95), while aDCHS1 organoids had an increased frequency of striatal Neurons (sNeurons, d=-1.00) (Fig. 4c,d, Supplementary Figs. 3e).

As these findings suggest that significant genotype-specific effects exist with regard to neural development and specialization, we validated these results through immunohistochemistry (IHC) staining of the aDCHS1 and hDCHS1 organoids. Given the altered balance between dRG and RG, which suggests a shift in the proportion between dorsal and striatal progenies, we decided to investigate the percentage of ventricular areas that would give rise to dorsal and striatal regions. In line with the scRNA-seq quantifications (Fig. 4e), the number of PAX6+ positive ventricular areas were less abundant in aDCHS1 than in hDCHS1 organoids (Fig. 4g). Furthermore, when present, these areas were also thinner (Supplementary Figs. 4a,f). Cells positive for markers for intermediate progenitors (TBR2+, Fig. 4i, Supplementary Figs. 4b) and deep layer neurons (TBR1+/CTIP2+, Supplementary Figs. 4c,d,h,i) were less abundant in aDCHS1 compared to the hDCHS1 organoids, while upper layer neurons (SATB2+) did not differ (Supplementary Figs. 4e,l). Conversely, we observed a higher proportion of MEIS2+ ventricular areas in aDCHS1 neural organoids at day 30 of LGE identity (Fig. 4h), which is known to give rise to the vast majority of striatal projection neurons (SPNs). Moreover, cells positive for striatal neuronal markers such as FOXP2 were more frequent in aDCHS1 than in hDCHS1 organoids (Fig. 4j). To validate this observed cell fate switch, we furthermore employed a model that predicts the dorsal or striatal molecular identity of cells based on their transcriptomic features by a Dorso-Striatal score (DS), derived from neural organoids and human cortex ^24,51^. The model indicated that aDCHS1 organoids have a greater proportion of cells with a striatal identity than hDCHS1 organoids, regardless of the organoids’ age (Fig. 4f, Supplementary Figs. 3g-i).

### EPHA4 binds DCHS1 with higher affinity in its ancestral form

To investigate how the presence of a sugar moiety at position 777 in DCHS1 modulates cell-cell communication and molecular interactions, we combined in silico predictions with experimental proteomic analyses (Fig. 5a). Specifically, we used the CellChat algorithm to analyze ligand-receptor interactions based on the previously described scRNA-seq data from neural organoids ^52^. Additionally, we performed mass spectrometric identification of co-precipitating proteins from organoids expressing hDCHS1 and aDCHS1, enabling a comprehensive assessment of potential interactors (Fig. 5a). CellChat inference suggested that several receptor-ligand pairs involved in neurodevelopmental processes might be differentially regulated in aDCHS1 organoids compared to hDCHS1 organoids (Fig. 5b). In particular, this analysis highlighted key neurodevelopmental receptors - including Ephrins, Semaforins, and Slit family - known to regulate neural cell fate, migration and corticogenesis ^53-55^. Guided by these predictions, we next sought to determine whether the glycosylation in aDCHS1 affects its binding partners in neural tissue. We purified tagged versions of aDCHS1 and hDCHS1, incubated them 60-day-old neural organoid lysates, and performed immunoprecipitation followed by mass spectrometric identification of co-precipitating proteins (Supplementary Figs. 5a). Integrating the proteomics and CellChat predictions, we identified EPHA4 as a prominent candidate (Fig. 5a-c, Supplementary Figs. 5b). Notably, EPHA4 has not been previously identified as a DCHS1 interactor, highlighting a potential novel interaction. Interestingly, we also detected shifts in the abundance of EPHA4 ligands, suggesting that glycosylation on aDCHS1 may alter EPHA4’s affinity for different ephrins. Specifically, EFNA5, EFNB1 and EFNB3 were differentially enriched in aDCHS1 pulldowns compared to hDCHS1 (Fig. 5c, Supplementary Figs. 5b). Although additional differentially expressed interactors were identified, the combined analyses strongly prioritized EPHA4 for further validation. Given the established role of Eph/ephrin signaling in regional progenitor specification ^56^, these changes could influence dorsal (cortical) versus ventral (striatal) identity by modulating progenitor positioning, adhesion, or signaling cues in the developing neural organoid. Collectively, these findings suggest that glycosylation at position 777 in DCHS1 might regulate its interaction with EPHA4, thereby influencing Eph/ephrin dynamics. Specifically, the differential co-immunoprecipitation of ephrin ligands in aDCHS1 versus hDCHS1 conditions points to shifts in Eph/ephrin-mediated boundary formation and progenitor allocation within the neural organoid.

**Fig. 5.**
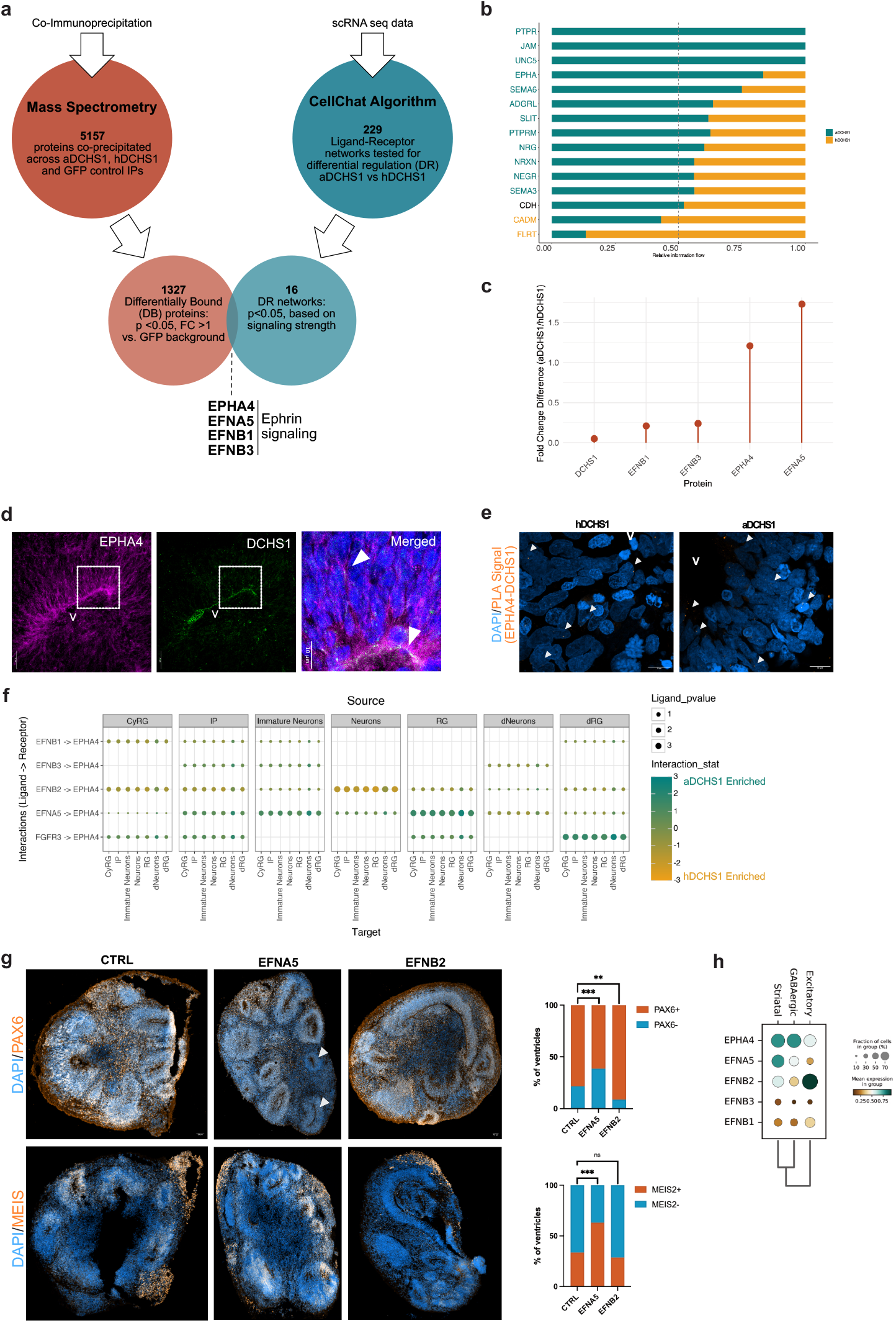
Alternative EPHA4 interaction with aDCHS1 may contribute to differences in cortico-striatal proportions. **(a)** Schematic representation of the selection process for ephrin signaling interactions. Co-immunoprecipitation mass spectrometry identified 5157 proteins, of which 1327 were differentially bound (DB; p < 0.05, FC > 1 vs. GFP). In parallel, CellChat analysis of scRNA-seq data tested 229 ligand–receptor networks, identifying 16 differentially regulated (DR; p < 0.05) interactions based on signaling strength. Ephrin signaling was selected as a key candidate pathway by intersecting both datasets, guiding further analysis of its role in aDCHS1 and hDCHS1 organoids. **(b)** Significant signaling pathways ranked based on differences in overall information flow within the inferred networks between aDCHS1 and hDCHS1. Pathways enriched in aDCHS1 are shown in teal, equally enriched pathways in black, and pathways enriched in hDCHS1 in yellow. **(c)** Fold-change differences in protein abundance from immunoprecipitation (IP) analysis between aDCHS1 and hDCHS1. **(d)** Representative fluorescence images of DAPI (blue), DCHS1 (green), and EPHA4 (magenta) in control 60-day-old neural organoids. *V* = ventricle; scale bars: 25 µm (individual channels) and 10 µm (merged image). **(e)** Representative fluorescence images of 60-day-old hDCHS1 and aDCHS1 neural organoids subjected to proximity ligation assay (PLA) to detect DCHS1-EPHA4 interactions. Nuclei were stained with DAPI. Images were acquired at 63× magnification. *V* = ventricle; scale bar: 10 µm. **(f)** Differential expression analysis of cell-cell communication events potentially deregulated between aDCHS1 and hDCHS1 related to the EPHA4 receptor. Dot color indicates enrichment in hDCHS1 (yellow) or aDCHS1 (teal). **(g)** Representative immunohistochemistry (IHC) images and quantification of PAX6+ and MEIS2+ cells in 30-day-old neural organoids derived from human iPSCs, treated with EFNA5, EFNB2, or left untreated. Statistical significance was assessed using a binomial test by comparing each treated condition to the untreated condition individually. For PAX6 (Batch=1), n = 4 organoids per condition; total number of ventricles: Untreated = 23, EFNA5 = 62, EFNB2 = 45. For MEIS2 (Batch=1), n = 4 organoids per condition; total number of ventricles: Untreated = 30, EFNA5 = 35, EFNB2 = 49. **(h)** Dot plot showing the expression of EPHA4, EFNA5, EFNB2, EFNB3 and EFNB1 genes in the Excitatory, Inhibitory and Striatal lineage. Dot color represents the average expression level, and dot size indicates the proportion of cells expressing each gene.

To test this hypothesis and further validate our immunoprecipitation results, we next sought to validate whether DCHS1 and EPHA4 interact in situ. We performed a proximity ligation assay (PLA) on 60-day-old neural organoid sections (Fig. 5e). As PLA detects only those signals that occur when two proteins are in close proximity (< 40 nm), these results provide robust confirmation of a direct or near-direct interaction between DCHS1 and EPHA4. Furthermore, immunohistochemical analyses (Fig. 5d) showed colocalization of DCHS1 and EPHA4 at distinct sites within apical radial glial cells in the ventricular zone, reinforcing the physiological relevance of the identified interaction and supporting a role for the DCHS1-EPHA4 axis in progenitor behavior during neurogenesis.

To further refine the CellChat-based predictions and to gain deeper insight into the differential ephrin ligand interactions initially identified in our immunoprecipitation experiments, we employed the ligand–receptor analysis tool LIANA on the same scRNA-seq datasets (Supplementary Figs. 5d). LIANA identifies potential cell–cell communication events by integrating multiple computational methods and evaluating differentially expressed ligand–receptor pairs. Among the most significant differentially expressed interactions, EPHA4 again emerged as a key receptor whose ligand–binding dynamics varied between aDCHS1 and hDCHS1 organoids (Fig. 5f, Supplementary Figs. 5d). Specifically, LIANA reinforced that EPHA4 shows altered binding patterns toward EFNB2 and EFNA5 in aDCHS1 compared to hDCHS1, suggesting a possible mechanism by which shifts in Eph/ephrin signaling could influence progenitor domain identity (i.e., cortical vs. striatal) in the developing neural organoid. Notably, while EPHA4 expression is relatively widespread across both dorsal and ventral progenitor domains, its physiological ligands, the ephrins EFNA5, EFNB1, EFNB2, exhibit more spatially patterned transcriptional expression (Fig. 5h, Supplementary Figs. 5c). Consistent with previous findings in the developing mouse cortex ^56,57.^ Given that ephrin ligands were suggested to drive region-specific neurodevelopmental outcomes by engaging EPHA4 in a context-dependent manner ^56,57,^ we next tested whether EFNA5 and EFNB2 could alter progenitor domain identities when administered exogenously to human-derived neural organoids. We added either EFNA5 or EFNB2 every 3 days for 15 days and performed immunohistochemical analyses on 30-day-old organoids to assess progenitor identity. Remarkably, EFNA5 treatment promoted the formation of LGE-like ventricles, evidenced by a marked increase in MEIS2+ progenitors, consistent with ventral striatal identity (Fig. 5g). Conversely, EFNB2 treatment resulted in an expansion of cortical-like ventricular zones, indicated by higher levels of PAX6+ progenitors (Fig. 5g).

These results confirm that EFNA5 and EFNB2 can differentially shape dorsal–ventral patterning in developing organoids, providing direct functional evidence for the involvement of these ligands in specifying cortical versus striatal lineages. When placed in the context of our earlier findings—namely, that ancestral (aDCHS1) and human (hDCHS1) variants of DCHS1 show distinct capacities to bind EPHA4 and modulate its interaction with ephrin ligands—this reinforces the concept of a dorsal–ventral “ephrin combinatorial code”. Perturbations in the DCHS1– EPHA4 axis may therefore drive shifts in cortico-striatal proportions, offering a mechanistic framework by which progenitor plasticity could be refined through evolutionary time. These observations advance our understanding of how the balance of cortical and striatal territories may have been modulated during primate and human brain evolution.

## Discussion

In this study, we determined how the loss of a single ancestral glycosylation site in human DCHS1 altered EPHA4-ephrin ligand signaling, thereby shifting cortico-striatal proportions that may have contributed to distinctive features of modern human brain evolution. Primates are known to have generally increased cortical volume compared to non-primate species; however, while the human telencephalon follows this trend, many of its substructures expand at different rates, with some actually showing relative decreases. Several studies have proposed that this reflects a reorganization of the cortex to support human-specific cognitive functions, most notably language. Consistent with these observations, we find that the striatum is among the brain structures that show a relative decrease in size.

This may facilitate the marked cerebral cortical expansion observed in humans, which could underlie the emergence of complex linguistic abilities^58-60^. Our findings using human, ancestralised, and chimpanzee iPSC-derived neural organoids provide experimental support for this early divergence in cortical-striatal proportions. Human organoids displayed an expanded pool of cortical progenitors and increased cortical compartment size, whereas chimpanzee organoids exhibited a larger striatal compartment relative to cortical structures. These results suggest that species-specific differences in neocortex-to-striatum proportions emerge early in development and are not solely a consequence of postnatal or activity-dependent mechanisms. Moreover, these differences align with fossil and endocast evidence, which suggests that cortical and subcortical reorganisations emerged progressively throughout hominid evolution, culminating in the pronounced neocortical expansion characteristic of *Homo sapiens* ^9,23,61^. In particular, recent fossil-based shape analyses of Neanderthal endocasts have revealed evolutionary changes in early brain development^62^, including differences in endocranial globularity in regions such as the prefrontal cortex and temporal lobes—reinforcing that telencephalic organization underwent significant modifications during the emergence of modern humans.

Building on the idea that specific hominid variants may have driven these structural divergences, we examined genes carrying modern human-specific missense variants. Previous work on introgressed Neanderthal SNVs has implicated *UBR4*, a ubiquitin ligase important for neurogenesis and migration in the developing brain, in shaping striatal regions such as the putamen ^9,63.^ Meanwhile, genes such as *FOXP2* and *ROBO1/2*, which are strongly associated with striatal function, lie within so-called “introgression deserts” - regions significantly depleted of archaic variants^64^. Our *in-silico* analyses show that around a quarter of modern human-specific missense variants converge on PTMs, particularly phosphorylation and glycosylation. As many mutations are present in genes expressed in the developing brain, these fine-scale proteomic modifications may significantly influence neurodevelopment, providing a mechanism by which subtle genetic changes lead to amplified developmental outcomes. This hypothesis is consistent with the general observation that evolutionary innovations often result from proteomic “tuning” rather than wholesale reorganization of transcriptional networks. Interestingly, although overall glycosylation complexity has increased in the hominid brain^64^, certain key glycosylation sites appear to have been lost in modern humans, suggesting that selective removal of certain glycans may also be advantageous.

In humans, DCHS1 missense mutations can be associated with brain malformations, suggesting that modulations of this gene may have profound effects on brain organization^43,65^. This modern human-specific change is functional because it stabilizes the protein, suggesting that loss of the ancestral glycan may affect DCHS1 conformation, adhesion properties, or both.

Our findings provide a mechanistic understanding of how subtle molecular modifications can drive large-scale shifts in brain organization across evolution. Single-cell RNA sequencing and immunohistochemical analyses revealed that the ancestral form of DCHS1 (aDCHS1) preferentially supports the expansion of ventral (striatal) progenitors, whereas the modern human variant (hDCHS1) favors increased cortical progenitor pools. This phenomenon mirrors broader evolutionary trends in primate brain development, where the neocortex of modern humans is comparatively expanded relative to that of other primates. A salient discovery of this study is the revelation that EPHA4, a hitherto unidentified interactor of DCHS1, demonstrates glycosylation-sensitive binding to ephrin ligands (EFNA5, EFNB2), thereby establishing a molecular link between DCHS1 function and regional progenitor identity. The capacity of exogenous EFNA5 and EFNB2 to modulate dorsal-ventral progenitor proportions in human organoids, which recapitulates findings from mouse experiments^56,57,66^, underscores the significance of Eph/ephrin signaling in cortical-striatal boundary formation. These findings imply that even modest biochemical alterations, such as the loss of a single glycosylation site, can calibrate receptor-ligand dynamics, resulting in substantial developmental reorientations. Nonetheless, while our work emphasizes the role of EPHA4 due to its established significance in brain regionalization, it is important to acknowledge that DCHS1 may engage with additional binding partners. Such interactors could independently or synergistically exert distinct developmental effects, representing compelling avenues for future research. Taken together, our results support a model in which a single amino acid substitution in DCHS1 - one that eliminates an ancestral glycosylation site - modulates the EPHA4-ephrin signaling axis to expand cortical territories at the expense of subcortical ones. More broadly, this work underscores how PTMs can function as evolutionary fine-tuning mechanisms, subtly adjusting developmental signaling pathways without requiring major transcriptional changes. The diverse biological roles of DCHS1, ranging from planar cell polarity in flies to progenitor specification in mammalian brains, illustrate how protocadherin family proteins have been repeatedly co-opted for new developmental tasks during evolution. Future research on other PTM-disrupting variants may reveal additional convergent or synergistic pathways underlying modern human brain organization and function.

### Limitations of the study

While our study has identified DCHS1 glycosylation as a key aspect of its evolutionary changes during brain development, some aspects require further investigation. Although we have shown that modern human DCHS1 has lost a glycosylation site - potentially contributing to changes in its stability and interaction with other proteins - the exact molecular mechanism remains partially understood. Glycosylation is generally stable once it reaches the plasma membrane, but studying how different types of N-glycan branching (microheterogeneity) might affect ancestral DCHS1 *in vitro* remains challenging.

One notable limitation of our study is the use of neural organoids to model mutations. While neural organoids represent a significant advancement in neuroscience research—providing a closer approximation to human brain development than traditional cell cultures and serving as a powerful tool to model SNVs from non-living relatives like Neanderthals—they do not fully capture the complexity and maturity of the human brain. In particular, neural organoids tend to represent early developmental stages and may not exhibit the same level of structural and functional organization as a fully developed brain. Moreover, although the striatum appears proportionally smaller in humans, its circuitry may have undergone significant reorganization. Future work employing more sophisticated or complementary systems will be necessary to investigate these later-developing features and fully understand how evolutionary shifts in connectivity and neurotransmission contribute to modern human brain function.

This SNV in DCHS1 provides substantial evidence for neuroanatomical changes in modern humans; however, it is merely one of many factors contributing to these changes. With at least 90 non-synonymous missense variants potentially influencing brain development— some of which may interact with each other—the complexity of these changes is likely to be higher than is currently appreciated. Additionally, the genetic background of modern humans may either amplify or diminish the effects of Neanderthal mutations, complicating direct comparisons between the two species. Given that some mutations affect multiple PTMs, their regulatory potential may be even greater. Joint analyses of *DCHS1, UBR4, FOXP2*, and *ROBO* - key genes in striatal development and introgression deserts - could clarify their collective role in shaping cognitive and structural differences between modern humans and other hominins. Recently, the groups of Giuseppe Testa and Cedric Boeckx investigated the synergistic roles of multiple genes that exhibit differences between ancestral and modern humans, particularly those crucial for brain development. Their findings underscore the significance of gene interactions, shedding light on how these interactions contribute to the complex process of brain evolution. In line with this, exploring the interplay of *DCHS1, UBR4, FOXP2*, and *ROBO* may reveal synergistic effects on brain evolution that single-gene studies might miss.

## Methods

### iPSCs lines and generation of aDCHS1 cell lines

For this study we used human 409-B2 hiPSC (female, Riken BioResource Center, catalog no. HPS0076), and chimpanzee SandraA ciPSC (female, Mora-Bermúdez et al. 2016 ^67^). For the Mora-Bermúdez et al. study, primate blood samples used to generate iPSCs were obtained by certified veterinarians during annual medical examinations or other necessary medical interventions, meaning that no invasive procedures were performed on primates for the sole purpose of our research project. The MPI EvA has an institutional permit for the transport of biological material derived from endangered species (DE216-08). iPSCs were cultured on Matrigel (Corning) coated plates (Thermo Fisher, Waltham, MA, USA) in mTesR1 basic medium supplemented with 1x mTesR1 supplement (STEMCELL Technologies, Vancouver, Canada) at 37 °C, 5% CO_2_, and ambient oxygen level. Passaging was done by Gentle Cell Dissociation Reagent (STEMCELL Technologies) treatment. All cell lines were tested for mycoplasma contamination before and after experiments.

Genome editing of DCHS1 in 409-B2 hiPSC to the ancestral state was done independently using Cas9D10A double nicking (gRNA_t1 GTGGACATCAGCATTGTGCC, gRNA_t2 GGGCACTGGGTTCTGCCTGT, Cas9D10A_donor: AAAAAACATACTGTAGTTGCTCAAATATGGGTGGTGTGGGGGTTCCAGGCACGATGCTGATGTTCACTCGGGCACTGGGTTCTGCCTGTAGGCCACCTCCGTCCTCAGCCCCGATCTCCAGCTGCACCACAGAATT), as well as using high-fidelity Cas9 nuclease (gRNA_t3 GCACAATGCTGATGTCCACT, Cas9_donor: GGCCCTTGGCCAGACGGGCCAATTCTGTGGTGCAGCTGGAGATCGGGGCTGAGGACGGAGGTGGCCTACAGGCAGAACCCAGTGCCCGAGTGAACATCAGCATTGTGCCTGGAACCCCCACACCACC). As both approaches used different gRNAs, shared phenotypes observed from ancestralized cells cannot be due to potential gRNA-dependent off-target effects. Electroporation was done as initially described by Riesenberg and Maricic ^68^ (GMO permit AZ 54-8452/26) using the B-16 program of the Nucleofector 2b device (Lonza) in cuvettes for 100 μl Human Stem Cell nucleofection buffer (Lonza, catalog no. VVPH-5022), containing 1 million cells, 100 pmol electroporation enhancer, 320 pmol gRNA (crRNA/tracR duplex), 200 pmol of single-stranded DNA donor, and 252 pmol CRISPR enzyme. We used recombinant Streptococcus pyogenes Cas9, Cas9-HiFi (R691A) and Cas9D10A proteins from Integrated DNA Technologies. Edited cells were plated in different wells, passaged, and then plated in a single cell dilution that gave rise to single cell-derived colonies.

Cells for analysis were dissociated using Accutase (SIGMA, A6964), pelleted, resuspended in 15 µl QuickExtract (Epicentre, QE0905T), and incubated at 65 °C for 10 min, 68 °C for 5 min, and finally 98 °C for 5 min. PCR of the editing site to confirm successful editing, as well as of heterozygous positions upstream and downstream to exclude loss-of-heterozygosity ^69^, was done in a T100 Thermal Cycler (Bio-Rad) using the KAPA2G Robust PCR Kit (SIGMA, KK5024) with supplied buffer B and 3 µl of cell extract in a total volume of 25 µl. The thermal cycling profile of the PCR was: 95 °C 3 min; 34 × (95° 15 s, 65 °C 15 s, 72 °C 15 s); 72 °C 60 s (DCHS1_forward: GGGTTGTGTGCCTGGACTAT, DCHS1_reverse: TTCCTCTCAGGGCTGTTGAC, rs2659870_forward: CCTGCACCTAAGAAGCTGGT, rs2659870_reverse GAGACTCAGGACCCAATGGA, rs72911031_forward: CTGCCCCAAAGCCATAATAA, rs72911031_reverse GCAACAGAGAGCCTGTCTCA). Sample-specific indices on P5 and P7 Illumina adapters were added in a second PCR reaction using Phusion HF MasterMix (Thermo Scientific, F-531L) and 0.3 µl of the first PCR product. The thermal cycling profile of the second PCR was: 98 °C 30 s; 25 × (98° 10 s, 58 °C 10 s, 72 °C 20 s); 72 °C 5 min. The indexed amplicons were purified using Solid Phase Reversible Immobilization (SPRI) beads ^70^. Double-indexed libraries were sequenced on a MiSeq (Illumina) giving paired-end sequences of 2 × 150 bp. After base calling using Bustard (Illumina), adapters were trimmed using leeHom ^71^, and sequences were analysed by SAMtools ^72^.

To exclude heterozygous large deletions that are invisible to amplicon PCR ^73^, we estimated the copy number of the target sequences by quantitative ddPCR. Primers were designed flanking the cut site and the probe was designed excluding edited sites. The gene *FOXP2* was used as a copy number reference. The ddPCR amplification was done in 1× ddPCR Supermix for probes (no dUTP, Bio-Rad, catalog no. 1863024), 0.2 μM primer and 0.2 μM probe for target and reference, together with 1 μl genomic DNA in QuickExtract DNA extraction solution (Lucigen, catalog no. QE09050). After droplet generation, the PCR reaction for DCHS1 (DCHS1_ddPCR_forward CACATCCTCTGGCACAGAAA, DCHS1_ddPCR_reverse GAGATCGGGGCTGAGGAC, DCHS1_ddPCR_probe 6FAM-TGGTGTGGGGGTTCCAGGCAC-BHQ_1) and FOXP2 (FOXP2_ddPCR_forward GCAACAGCAATTGGCAGC, FOXP2_ddPCR_reverse CAGCGATTGGACAGGAAGTG, FOXP2_ddPCR_probe HEX-AGCAGCAGCAGCATCTGCTCAGCCT-BHQ_1) was run for 5 min at 95 °C, followed by 42 cycles of 35 s at 95 °C (at a ramp rate of 1.5 °C s−1) and 65 s at 60 °C (at a ramp rate of 1.5 °C s−1) and 5 min at 98 °C. Droplets were read in a QX200 Droplet reader (Bio-Rad) and allele copy numbers were determined relative to a different fluorophore for the FOXP2 reference and unedited control.

Karyotyping by trypsin induced Giemsa staining to confirm a healthy diploid karyotype of studied cellular clones were carried out according to international quality guidelines (ISCN 2016: An International System for Human Cytogenetic Nomenclature ^74^) by the ‘Sächsischer Inkubator für klinische Translation’ (Leipzig, Germany).

### Neural organoids generation

Neural organoids were generated following Lancaster et al. 2014 directions with some small changes ^75^. iPSCs were dissociated into single cells StemPro Accutase Cell Dissociation Reagent (A1110501, Life Technologies) and plated in the concentration of 9000 single iPSCs/well into low-attachment 96-well tissue culture plates in hES medium (DMEM/F12GlutaMAX supplemented with 20% Knockout Serum Replacement, 3% ES-grade FBS, 1% nonessential amino acids, 0.1 mM 2-mercaptoethanol, 4 ng/mL bFGF, and 50 µM Rock inhibitor Y27632) for 6d in order to form embryoid bodies (EBs). Rock inhibitor Y27632 and bFGF were removed on the 4th day. On day 6, EBs were transferred into low-attachment 24-well plates in NIM medium (DMEM/F12GlutaMAX supplemented with 1:100 N2 supplement, 1% nonessential amino acids and 5 µg/mL Heparin) and cultured for additional 6d. On day 12, EBs were embedded in Matrigel (Corning, 354234) drops and then they were transferred in 10 cm tissue culture plates in NDM-A medium (DMEM/F12GlutaMAX and Neurobasal in ratio 1:1 supplemented with 1:100 N2 supplement 1:100 B27 without vitamin A, 0.5% nonessential amino acids, insulin 2.5 µg/mL, 1:100 Antibiotic-Antimycotic, and 50 µM 2-mercaptoethanol) in order to form neural organoids (NOs). 4 days after Matrigel embedding, NOs were transferred into an orbital shaker and cultured until electroporation in NDM+A medium (DMEM/F12GlutaMAX and Neurobasal in ratio 1:1 supplemented with 1:100 N2 supplement 1:100 B27 with vitamin A, 0.5% nonessential amino acids, insulin 2.5 µg/mL, 1:100 antibiotic-antimycotic, and 50 µM 2-mercaptoethanol). During the whole period of neural organoids generation, cells were kept at 37°C, 5% CO2, and ambient oxygen level with medium changes every other day. After transferring the NOs onto the shaker, the medium was changed twice per week. NOs were cultured up to 30, and 60 days as indicated.

In the case of chimp iPSCs, or when human iPSCs were compared to chimp lines, neural organoids were generated with a few modifications. 24 hours prior EB generation, iPSCs were treated with 1% DMSO in mTeSR1 media. iPSCs were dissociated into single cells StemPro Accutase Cell Dissociation Reagent (A1110501, Life Technologies) and plated in the concentration of 2000 single iPSCs/well into low-attachment 96-well tissue culture plates in mTeSR medium supplementer with Rock inhibitor (50 µM final concentration). After 48 hours, media was replaced with Forebrain medium 1 (DMEM/F12 supplemented with 20% Knockout Serum Replacement, 1% nonessential amino acids, 1x GlutaMAX supplement, 0.1 mM 2-mercaptoethanol and 1x antibiotic-antimycotic) supplemented with 100 ng/ml Noggin (StemCell, Cat no. 78060). Media was replaced any second day and on day 5, media was replaced with NIM medium supplemented with 100 ng/ml Noggin. On day 7, EBs were embedded in Matrigel Cookies as explained in Qian et al., 2018 (76). On day 9, media was replaced to IDM-A (DMEM/F12-GlutaMAX and Neurobasal in ratio 1:1 supplemented with 1:200 N2 supplement 1:50 B27 without vitamin A, 0.5% nonessential amino acids, insulin 2.5 µg/mL, 1:100 Antibiotic-Antimycotic, and 50 µM 2-mercaptoethanol) and changed every second day. On day 14, Matrigel cookie was removed and media was replaced by IDM+A (DMEM/F12-GlutaMAX and Neurobasal in ratio 1:1 supplemented with 1:200 N2 supplement 1:50 B27 with vitamin A, 0.5% nonessential amino acids, insulin 1 µg/mL, 1:100 antibiotic-antimycotic, and 50 µM 2-mercaptoethanol, 1 mg/ml NaHCO_3_) and organoids were transferred onto the orbital shaker and medium was replaced twice per week. Human brain organoids were treated with either recombinant EFNA5 (MCE, Cat. No.: HY-P70379) or EFNB2 (MCE, Cat. No.: HY-P77645) protein to study their effects on organoid development. The treatments were administered starting on day 5 of differentiation and continued until day 20. Both EFNA5 and EFNB2 were used at a final concentration of 0.5 µg/ml, with the treatment being included during every media change. The organoids were cultured under standard conditions, with media replenishment and treatment applied as described. NOs were fixed using 4% PFA for 1 h at 4 °C, cryopreserved with 30% sucrose for 24 hours and stored at −20 °C. For immunofluorescence, 20 µm cryosections were prepared. For each experiment, many independent ventricles per NO from at least 3 different NOs generated in 2–3 independent batches were analyzed.

### PTMs prediction analyses

Prediction of PTMs in genes linked to Neandertal mutations was performed by processing the full-length sequences of the human and Neanderthal versions using MusiteDeep ^36-38^. Proteins were screened for phosphorylation, glycosylation, ubiquitination, SUMOylation, acetylation, methylation, pyrrolidone carboxylic acid, palmitoylation, and hydroxylation. For glycosylation prediction, both aDCHS1 and hDCHS1 sequences were also uploaded to NetNGlyc-1.066 ^76^, a tool specializing in N-Glycosylation. In addition to automated predictions, PTMs were manually checked for their predicted cellular location to ensure they were consistent with known biological context (e.g., glycosylation being restricted to extracellular regions).

### DCHS1 plasmid generation

hDCHS1-TurboGFP plasmid was generated according to Di Matteo et al 2025 ^65^. aDCHS1-TurboGFP was generated using the hDCHS1-TurboGFP as template with the Q5^®^ Site-Directed Mutagenesis Kit (E0554), following the provider’s instructions. Primers for the codon mutations (gac to aac) were generated using the NEBaseChanger Tool (v2.5.0).

### Production of DCHS1 protein and MS analysis

To overexpression and purification of either hDCHS1 or aDCHS1, hDCHS1-GFP or aDCHS1-GFP plasmids were transiently expressed in HEK293 cells using PEI STAR™ transfection reagent (7854/100). 10 µg of plasmid was incubated with 20 µg of PEI for every 2 × 10^6^ cells, and 48 hours post-transfection, cells were harvested. Cells were resuspended in PBS and collected via centrifugation at 300 g for 3 minutes. Cell pellet was then lysed in ice cold IP lysis buffer (10 mM Tris/Cl pH 7.5, 150 mM NaCl, 0.5 mM EDTA, 0.5 % Nonidet™ P40 Substitute) and incubated on ice for 30 minutes with occasional pipetting. Lysates were clarified by centrifugation at 4°C for 10 mins at 16000 g and supernatant was collected in a new tube. To immunoprecipitated DCHS1, 25 µl TurboGFP-Trap Magnetic Agarose (Proteintech) beads were used every 200 µl of lysate following manufacturer’s instructions. Lysates were incubated with the beads for 1 hour at 4°C in an end-over-end rotator. Beads were then washed 3 times in wash buffer (10 mM Tris/Cl pH 7.5, 150 mM NaCl, 0.05 % Nonidet™ P40 Substitute, 0.5 mM EDTA) and in the last step beads were moved to a new tube to avoid any contamination of unbound proteins. Finally, DCHS1-TurboGFP were eluted from the beads via addition of 100 µl of acidic elution buffer (200 mM glycine pH 2.5) and eluates were neutralized with 5 µl of neutralization buffer (1 M Tris pH 10.4). To 100 uL of DCHS1 pulldown 3.2 uL 1 M HCl, 4 uL 250 mM pH neutralized TCEP, and 100 ng ProAlanase were added. The mixture was incubated for 2 h at 37°C and then directly cleaned up using 5 uL C18 cartridges on the Agilent AssayMap according to the manufacturer’s instructions. Peptides were dried in a SpeedVac and reconstituted in 0.1 % formic acid in water. LC-MS data acquisition was performed on an Orbitrap Eclipse equipped with a FAIMS Pro and an Easy Spray source coupled to a Vanquish Neo LC. Peptides were separated on a uPAC HT Neo column heated to 50 C coupled to a 30 um stainless steel emitter at a flow rate of 2.5 uL/min using a gradient from 1 to 5 % B within 0.5 min, 5 to 40 % over 22 min, 40 to 100 % B over 1 min, followed by a wash with 100 % B for 2 min with solvent A consisting of 0.1 % formic acid in water and solvent B consisting of 0.1 % formic acid in acetonitrile. The column was equilibrated with 1% B and a column equilibration factor of 1.5. Sample was directly injected onto the column at a pressure of 400 bar. The MS was operated in data dependent acquisition mode with a precursor scan in the Orbitrap at a resolution of 60k followed by MS2 acquisition of the top 15 most abundant precursors in the Orbitrap with a resolution of 30k. The precursor isolation width was set to 1.4 Th with an AGC target of 100 % and a max injection time of 54 ms using EThcD activation with calibrated charge dependent ETD parameters and a supplemental activation collision energy of 20 %. Raw DDA data was analyzed with Fragpipe v21.1, MSFragger 4.0, IonQuant 1.10.12, Philosoper 5.1.0, Python 3.9.13, and EasyPQP 0.1.42 using a fasta file consisting of the *Homo sapiens* and *Homo neanderthalensis* DCHS1 protein isoforms.

### Interactome analysis

Immunoprecipitation of DCHS1 was performed as described above. DCHS1-TurboGFP or empty TurboGFP plasmid on magnetic beads was subjected to 1 mg of neural organoids lysate prepared as follows. 60 days old control neural organoids were used for this experiment. 3 organoids per condition were lysed in IP lysis buffer and incubated on ice for 30 minutes with occasional pipetting. Lysates were clarified by centrifugation at 4°C for 10 mins at 16000 g and supernatant was collected in a new tube. DCHS1-TurboGFP-beads were incubated with the lysate for 1 hour at 4C in an end-over-end rotator. Beads were then washed 3 times in wash buffer (10 mM Tris/Cl pH 7.5, 150 mM NaCl, 0.05 % Nonidet™ P40 Substitute, 0.5 mM EDTA) and in the last step beads were moved to a new tube to avoid any contamination of unbound proteins. Finally, samples were eluted from the beads via addition of 200 µl of acidic elution buffer (200 mM glycine pH 2.5) and eluates were neutralized with 10 µl of neutralization buffer (1 M Tris pH 10.4). Proteins were denatured by addition of 100 μL of SDC buffer containing 1% sodium deoxycholate (SDC, Sigma-Aldrich), 40 mM 2-chloroacetamide (CAA, Sigma-Aldrich), 10 mM tris(2-carboxyethyl)phosphine (TCEP, Thermo Fisher Scientific), and 100 mM Tris, pH 8.0. The mixture was incubated for 30 minutes at 37°C. After incubation, proteins were digested overnight at 37°C with 0.5 µg of trypsin (Promega). Following digestion, the peptide solution was acidified with trifluoroacetic acid (TFA, Merck) to a final concentration of 1%. Desalting was then performed using SCX stage tips. A total amount of 200 ng of peptide material was loaded on Evotips (Evotip Pure, Evosep). Peptides were eluted from Evotips onto a 15 cm PepSep C18 column (15 cm x 15 cm, 1.5 µm, Bruker Daltonics) using the Evosep One HPLC system. The column was heated to 50°C, and peptides were separated using the 30 SPD method. Data acquisition was performed on a timsTOF Pro mass spectrometer using timsControl software. The instrument operated in data-independent (DIA) PASEF mode with a mass scan range of 100–1700 m/z and an ion mobility range of 1/K0 = 0.70 Vs cm^−2^ to 1.30 Vs cm^−2^. Equal ion accumulation and ramp time of 100 ms each were set in the dual TIMS analyzer, with a spectral rate of 9.52 Hz. DIA-PASEF scans were acquired in the mass range of 350.2–1199.9 Da, with an ion mobility range of 1/K0 = 0.70 Vs cm^−2^ to 1.30 Vs cm^−2^. The collision energy was ramped linearly from 45 eV at 1/K0 = 1.30 Vs cm^−2^ to 27 eV at 1/K0 = 0.85 Vs cm^−2^. A total of 42 DIA-PASEF windows were distributed to one TIMS scan each, with switching precursor isolation windows, resulting in an estimated cycle time of 2.21 seconds. Raw data were processed using Spectronaut 18.0 in directDIA+ (library-free) mode. The peak list was searched against a predicted human database from Uniprot (SwissProt and TrEMBL, downloaded in 2023) and custom FASTA sequences of bait proteins. Cysteine carbamidomethylation was set as a static modification, while methionine oxidation and N-terminal acetylation were set as variable modifications. Protein quantification across samples was performed using label-free quantification (MaxLFQ) at the MS2 level. Data processing was performed using the DEP (Differential Expression for Proteomics) package for statistical analysis, and the dplyr package for data manipulation. The protein identification data was first cleaned by removing irrelevant columns, excluding keratin proteins, and handling missing values. The readxl package was used to import the raw data, while the BiocManager package ensured the necessary dependencies were installed. Protein quantification was normalized using variance stabilization normalization (VSN) using functions from the DEP package. Differential expression analysis was carried out using linear models with empirical Bayes statistics, comparing the experimental conditions to controls. Statistical significance was determined using a threshold of p-value < 0.05 and log2 fold change > 0.5. Volcano plots, heatmaps, and principal component analysis (PCA) were generated using ggplot2 and other related visualization tools. To identify functional enrichment, gene ontology (GO) enrichment analysis was performed on the upregulated proteins using the gprofiler2 package, which provides analysis of GO terms, biological processes, and cellular components.

### CTSA assay

DCHS1-TurboGFP (either the ancestral or human version) was overexpressed in HEK293 cells as described above in T75 flasks. Following cell harvest, the cells were washed twice in PBS by gentle pipetting and centrifugation at 300g for 4 minutes each time. The washed cells were then resuspended in 0.9 ml of PBS containing a Protease Inhibitor Cocktail (PIC), and the suspension was divided into 8 PCR tubes (50 μl per tube). Each sample underwent a single heat shock cycle, consisting of 6 minutes at 40°C, followed by 6 minutes of cooling down to 25°C. Subsequently, NP-40 was added to each tube to a final concentration of 0.3%. The samples were then subjected to 3 rounds of snap-freezing and thawing: 5 minutes on dry ice followed by 5 minutes at room temperature, to facilitate cell lysis. After the cycles, the lysates were clarified by centrifugation at 15,000g for 30 minutes to remove denatured proteins. The soluble fraction was collected and resuspended in Laemmli buffer for further analysis.

### Western blotting

50 µl of samples from the CTSA assay were loaded on a 6% SDS-PAGE polyacrylamide gel run at 100V in the running buffer (25 mM Tris, 192 mM glycine, 0.1% SDS). After separation, proteins were transferred to PVDF membrane using a wet-transfer system at 100V for 120 mins in transfer buffer (10% MeOH, 0.03% SDS, 25 mM Tris, 192 mM glycine). Membranes were blocked in 5% BSA in TBS-T (0.01%) for 1 hour at room temperature and incubated with primary antibody diluted in 5% BSA in TBS-T overnight at 4°C with gentle shaking. After washing three times with TBS-T, membranes were incubated with LICOR secondary antibody for 1 hour at room temperature. Following three washes in TBS-T and one last wash in TBS, protein bands were detected using the LICOR Imaging system. Bands were quantified using ImageJ software.

### scRNA seq sample preparation and analysis

Single-cell dissociation was performed on five 30 and 60 days old neural organoids randomly selected for each condition. Single cells were dissociated using StemPro Accutase Cell Dissociation Reagent (Life Technologies), filtered through 30 uM and 20 uM filters (Miltenyi Biotec) and cleaned of debris using a Percoll (Sigma, P1644) gradient. Single cells were resuspended in ice-cold Phosphate-Buffered Saline (PBS) supplemented with 0.04% Bovine Serum Albumin at a concentration of 1000 cells per ul. Single cells were loaded onto a Chromium Next GEM Single Cell 3′ chip (Chromium Next GEM Chip G Single Cell Kit, 16 rxns 10XGenomics PN-1000127) with the Chromium Next GEM Single Cell 3′ GEM, Library & Gel Bead Kit v3.1 (Chromium Next GEM Single Cell 3′ GEM, Library & Gel Bead Kit v3.1, 4 rxns 10xGenomics PN-1000128) and cDNA libraries were generated with the Single Index Kit T Set A, 96 rxns (10xGenomics PN-1000213) according to the manufacturer’s instructions ^77^. Libraries were sequenced using Illumina NovaSeq6000 in 28/8/91bp mode (SP flowcell). Quality control and UMI counting and aggregation of samples were performed using the Cell Ranger software (version 6.0.0) using as a reference genome GRCh38-1.2.0. Downstream analysis was performed using the Python package Scanpy (version 1.9.6) following Huemos et al. guidelines ^78,79^. Low-quality cells were filtered using median absolute deviation (MAD), and were marked as outlier cells that differ by 5 MADs. Ambient RNA correction was applied using the SoupX tool ^80^. Following the correction, quality control (QC) filtering was performed on the single-cell RNA sequencing data using the scanpy package in Python. Cells with fewer than 1000 counts were filtered out, and cells with more than 40,000 counts were removed. Additionally, cells with fewer than 200 genes were excluded from the dataset. After each filtering step, the number of remaining cells was printed for tracking purposes. Once the QC filtering was complete, doublet detection was performed using the scDblFinder package (version 1.16.0) to identify and exclude doublets from the dataset ^81^. Moreover, we removed cells that were annotated as ‘doublets’ using the scDblFinder package (version 1.16.0). Normalisation of raw counts was performed using shifted logarithm transformation. The top 30 principal components were used for clustering (resolution of 0.5) using the Leiden algorithm from Scanpy ^82^. Clusters were grouped based on the expression of known marker genes and differentially expressed genes using the Wilcoxon rank-sum test. Moreover, we confirmed our manual annotation using CellTypist to unbiasedly annotate cells and map our cells to cell types from the first-trimester developing human brain ^83^. Cells which were predicted to be of ‘brain fibroblast’ identity were excluded as well as cells that were strongly predicted to have diencephalic origin. The 60 day old organoids’ dataset was integrated using Scanorama (version 1.7.4) and the standard workflows for visualization with UMAP and clustering were then used ^84^. Cell-cell communication analysis was performed using the CellChat package (version 2.1.2) in R ^52^. Two datasets, aDCHS1 and hDCHS1, were loaded and merged into a single CellChat object. The analysis began by comparing ligand-receptor interactions between the datasets, with visualizations generated to show differences in interaction networks. Heatmaps for signaling pathways were also produced, illustrating the interactions in both merged and individual datasets. Differential interactions were visualized using netVisual functions, comparing both count-based and weight-based measures. To investigate the signaling roles of specific cell types, scatter plots were generated using the netAnalysis functions, which examined signaling changes in early neural progenitor cells and striatal projection neurons. Additionally, functional and structural comparisons of the signaling networks were performed using manifold learning and clustering methods. Finally, network similarity was ranked, and visualizations were generated to compare the functional and structural similarities between the signaling networks, using custom color schemes to highlight differences between the cell groups. For the targeted cell-cell communication analysis, we employed the Liana package ^85,86.^ The AnnData object containing the processed single-cell RNA-sequencing data was used as input. Ligand-receptor interactions were inferred using multiple methods integrated within liana, including CellPhoneDB, CellChat, NATMI, SingleCellSignalR, and Connectome.

Pseudobulk analysis was performed on a subset of cells, specifically the radial glial (RG) cell type. Cells were first filtered based on their cell type and then grouped by the type column in the obs attribute of the AnnData object. For each group, pseudo-replicates were generated by randomly splitting the dataset into 6 subsets. Raw count data was used for the analysis, and each pseudo-replicate was created by summing the counts across cells within the subset. For each sample, an AnnData object was created for each pseudo-replicate, which was then added to a list of pseudobulk samples. These replicates were concatenated to form a combined pseudobulk dataset. The resulting dataset was analyzed using DESeq2 to perform differential expression analysis between the two conditions (aDCHS1 and hDCHS1) ^87^. The DESeq2 analysis was performed with the replicate and type columns as design factors. Principal component analysis (PCA) was conducted to visualize the variation in the data, and differential expression results were summarized and ranked by statistical significance and visualised using ggplot2 in Rstudio.

All computational analyses were performed using Python (version 3.9.18) in a Jupyter Notebook environment (jupyter_client 8.6.2, jupyter_core 5.7.2). Preprocessing and downstream analysis of single-cell RNA sequencing data were conducted using Scanpy (1.9.5), with visualization facilitated by matplotlib (3.8.2) and seaborn (0.13.2). Data structures were handled using pandas (2.2.3) and numpy (1.26.4). Cell type annotation was validated using CellTypist (1.6.3), and batch correction and dataset integration across organoid conditions were performed using Scanorama (1.7.4). The session environment was documented using session_info (1.0.0).

### Immunohistochemistry

Sections were equilibrated at room temperature and re-hydrated using DPBS for 10 minutes. Blocking was performed on section using 0.2% Tween, 5% Normal Goat Serum and 150 mM Glycine in DPBS for 1 hour at room temperature. Immunostaining on section was performed by incubating primary antibodies in blocking solution (made of 0.2% Tween, 5% Normal Goat Serum in DPBS) overnight at 4°C in a humidified chamber. Excess of primary antibody was removed by washing three times in PBS supplemented with 0.1% Tween. Nuclear staining (0.5 µg/mL 4,6-diamidino-2-phenylindole (DAPI, Sigma Aldrich)) and secondary antibodies were added at a concentration of 1:500 [vol/vol] in blocking buffer for 1 hour at room temperature. Sections were then washed 3 times in PBS supplemented with 0.1% Tween and one last time in PBS only, and finally mounted using Aqua-Poly Mount solution. For Proximity Ligation assay, the Duolink In Situ Red Starter Kit Mouse/Rabbit was used, following the manufacturer’s guidelines.

### Proximity Ligation Assay (PLA)

PLA was performed to detect protein-protein interactions in fixed cells or tissues using the Duo92101 PLA kit (Sigma-Aldrich). Primary antibodies against EPHA4 and DCHS1 were diluted in blocking buffer and incubated with the cells overnight at 4°C. Negative control reactions were performed by omitting either of the two antibodies. After washing with PBS, the PLA probes (designed to bind to the primary antibodies) were added according to the manufacturer’s protocol. The primary antibodies were then detected by the addition of oligonucleotide-conjugated secondary antibodies (PLA probes), which are complementary to the specific sequences present on the primary antibodies.

For ligation, the samples were incubated with a ligase solution that enables the proximity of two PLA probes within a 40 nm distance to form a circular DNA structure. The resulting DNA circle was amplified by rolling circle amplification (RCA) to produce fluorescent signals detectable by microscopy. The fluorescent signals indicate the proximity between the proteins of interest, providing evidence of protein-protein interaction. Cells were counterstained with DAPI to visualize nuclei, and images were acquired using a fluorescence microscope.

### Other bioinformatic analyses

DCHS1 sequences of the species of interest were aligned using Clustal Omega ^88^. The alignment was performed using the command-line version of Clustal Omega, and the resulting alignment was saved in FASTA format. The aligned sequences were then visualized using the ggmsa package in R. The sequence logo was colored according to the chemical properties of the amino acids using the “Chemistry_AA” color scheme, providing insights into the conservation of amino acids across the sequences in this region.

Bioinformatic analyses on the developing human brain at the first trimester were conducted using data extracted from the Cell Atlas of the Developing Brain ^24^. Expression matrices were downloaded from the Linnarsson GitHub page and processed using the scanpy library. To isolate specific cell classes and regions, the dataset was filtered to include only the desired cell types: Neuroblast, Radial Glia, Neuronal IPC, Neuroblast and Neuron. The dataset was further filtered to include cells from the regions Telencephalon, Forebrain, Brain, and Head. Unwanted regions, such as Diencephalon, Midbrain, Cerebellum, Medulla, Pons, and Hindbrain, were excluded using the Region attribute. Similarly, unwanted subregions, including Subcortex, Head, Hippocampus, and Brain, were excluded using the Subregion attribute. Cells with zero counts were filtered out using a minimum count threshold of 1, and normalization was performed by scaling the total counts per cell to 10,000 and applying a logarithmic transformation. Highly variable genes (HVGs) were selected using the Seurat v3 method, retaining the top 5,000 genes. Principal component analysis (PCA) was performed with 40 principal components, and the nearest neighbor graph was computed with 15 neighbors. UMAP dimensionality reduction was performed for visualization. Cells expressing *FOXG1* and *EMX1* were annotated as ‘Excitatory’, and cells expressing *DLX2* and *FOXG1*, but not *EMX1*, were annotated as ‘Inhibitory’. Cells from the Striatum were annotated based on the Subregion attribute and identified using a list of striatal markers, including *ISL1, ASCL1, DLX1*, and *EBF1*. The data were filtered to focus on ‘Excitatory’, ‘Inhibitory’, and ‘Striatum’ cells, and clustering was repeated as necessary.

For the dorso-ventral model, the data was split into training and testing sets. Feature selection was performed to identify the top 100 most significant genes. The selected genes were then used to train a Random Forest Classifier, employing stratified k-fold cross-validation to ensure robust model evaluation. The classifier’s performance was assessed using cross-validation scores and test set accuracy, ensuring the reliability of the selected features and the model’s predictive power.

Cohen’s d was computed to evaluate the effect size between two groups, hDCHS1 and aDCHS1, for each cluster annotation. The data consisted of a DataFrame containing the proportions of each cluster per sample, with the samples classified into two groups based on the sample names. A new column, type, was added to the dataset to distinguish between the two groups, which were used for grouping the data. The dataset was filtered to include only the relevant columns for analysis. For each cluster annotation, Cohen’s d was calculated by first determining the mean values for each group (denoted as M1 for hDCHS1 and M2 for aDCHS1). The standard deviations for each group (S1 for hDCHS1 and S2 for aDCHS1) were also computed. The pooled standard deviation (denoted as S_pooled) was calculated using the formula:

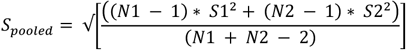

where N1 and N2 are the sample sizes for each group. Cohen’s d was then computed by subtracting the mean of the second group (M2) from the mean of the first group (M1), and dividing by the pooled standard deviation (S_pooled). If the pooled standard deviation was zero (indicating no variability in both groups), Cohen’s d was set to NaN to avoid division by zero. The computed Cohen’s d values for each cluster annotation were then compiled into a DataFrame, which was displayed in the Jupyter notebook. The analysis was performed using the Python libraries NumPy for numerical operations, including mean and standard deviation calculations, and Pandas for data manipulation.

The data for scRNA seq analysis of missense genes in the primate brain organoids was obtained from the paper Kanton et al., 2019 ^2^, and is available in the E-MTAB-7552 study hosted on the EBI biostudies platform (https://www.ebi.ac.uk/biostudies/arrayexpress/studies/E-MTAB-7552). This dataset includes single-cell RNA sequencing data for human, macaque, and chimp cells. The data files were downloaded using a bash script that accessed the EBI FTP server. The following files were retrieved: sparse matrices for cell counts (*.mtx), gene names (genes_consensus.txt), and metadata for each species (metadata_human_cells.tsv, metadata_macaque_cells.tsv, metadata_chimp_cells.tsv). The downloaded data was loaded into Python using the scanpy package. A custom function was used to read the sparse matrix files, load the gene names and metadata, and create an AnnData object for each species. The data was then concatenated into one AnnData object with an additional column (species) to distinguish between the three species (human, macaque, and chimp). The species column was added to the obs attribute to label each cell by its species. Principal component analysis (PCA) was performed on the combined dataset without filtering for highly variable genes (HVGs). A neighbor graph was computed using 15 nearest neighbors and 40 principal components (PCs). Uniform manifold approximation and projection (UMAP) was then performed for dimensionality reduction. For visualization, dotplots of gene expression across species were generated, focusing on a filtered list of genes and grouping by predicted cell type.

#### Prediction and Structural Modeling of DCHS1

For the generation of the DCHS1 structure, the AlphaFold3 server was used to predict the structure of the first 25 cadherin domains (amino acids 43-2706) of DCHS1 ^89^. Additionally, the glycan NAG(NAG(MAN(MAN(MAN)(MAN(MAN)(MAN))))) was modeled as an oligomannose on Asp 777 of aDCHS1. Molecular graphics and analyses were performed using UCSF ChimeraX, developed by the Resource for Biocomputing, Visualization, and Informatics at the University of California, San Francisco, with support from the National Institutes of Health (R01-GM129325) and the Office of Cyber Infrastructure and Computational Biology, National Institute of Allergy and Infectious Diseases.

#### Gene-Disease Association Analysis

Gene-disease association data was retrieved from the DisGeNET database using the DisGeNET API ^25,26^ .A list of candidate genes of interest was prepared using the genes linked to missense variants in Neanderthals. The gene symbols were URL-encoded to create a query string for the API. The request was made to the DisGeNET API endpoint (https://api.disgenet.com/api/v1/gda/summary), specifying the selected genes and filtering for neurological disorders (dis_class_list=“C10”). The response from the API, in JSON format, was parsed and processed using the httr and jsonlite packages in R. The relevant data, including gene symbols and associated disease names, was extracted and stored in a data frame. The gene-disease associations were then visualized using a heatmap, created by converting the data into a binary matrix indicating the presence (1) or absence (0) of associations for each gene-disease pair. The matrix was reshaped for use with ggplot2, and the heatmap was plotted to highlight the strength and frequency of gene-disease associations.

## Acknowledgements

We are deeply grateful to Svante Pääbo for inspiring this project and for his insightful discussions and critical comments to the manuscript, as well as to Denis Jabaudon for his critical reading and valuable suggestions, and Cedric Boeckx and Giuseppe Testa for enriching discussions. We thank Niklas Kroner-Weigl, Sarah Zund, Melina Vaki, Celeste Vervoort, and Maik Kodel for their invaluable technical support throughout the project. We also extend our appreciation to S. Robertson for generously providing the DCHS1GFP plasmid, and to the Mass Spectrometry Core Facility at MPI Biochemistry (RRID:SCR_025745) and the Sequencing Facility at MPI Molecular Genetics for their contributions. This work was supported by ERA-Net E-Rare (HETEROMICS | 01GM1914, S.C and A.F.E.), the European Union (ERC Consolidator Grant, ExoDevo | 101043959, S.C. and A.F.E.), the Human Frontiers Science Program (M.V.P.), the Alexander von Humboldt Foundation (M.V.P.), and the Max Planck Society. We also thank members of the Cappello, Nguyen, Baulac, Jabaudon, and Baffet labs for their collaborative spirit. We used ChatGPT (OpenAI) to assist with coding, data visualization, and workflow optimization in Python and R.

## Author information

S.C. and S.R. conceived the study; S.R. generated the ancestral cell lines; M.V.P., A.A.M., A.F.E.,G.B. and L.F. performed experiments; T.H. and M.M. performed mass spectrometry, M.V.P. analyzed data and M.V.P. and S.C. interpreted the data and wrote the manuscript with input from all authors.

