## Supplementary Figures for "DCHS1 Modulates Forebrain Proportions in Modern Humans via a Glycosylation Change"

### Supplementary Figure legends

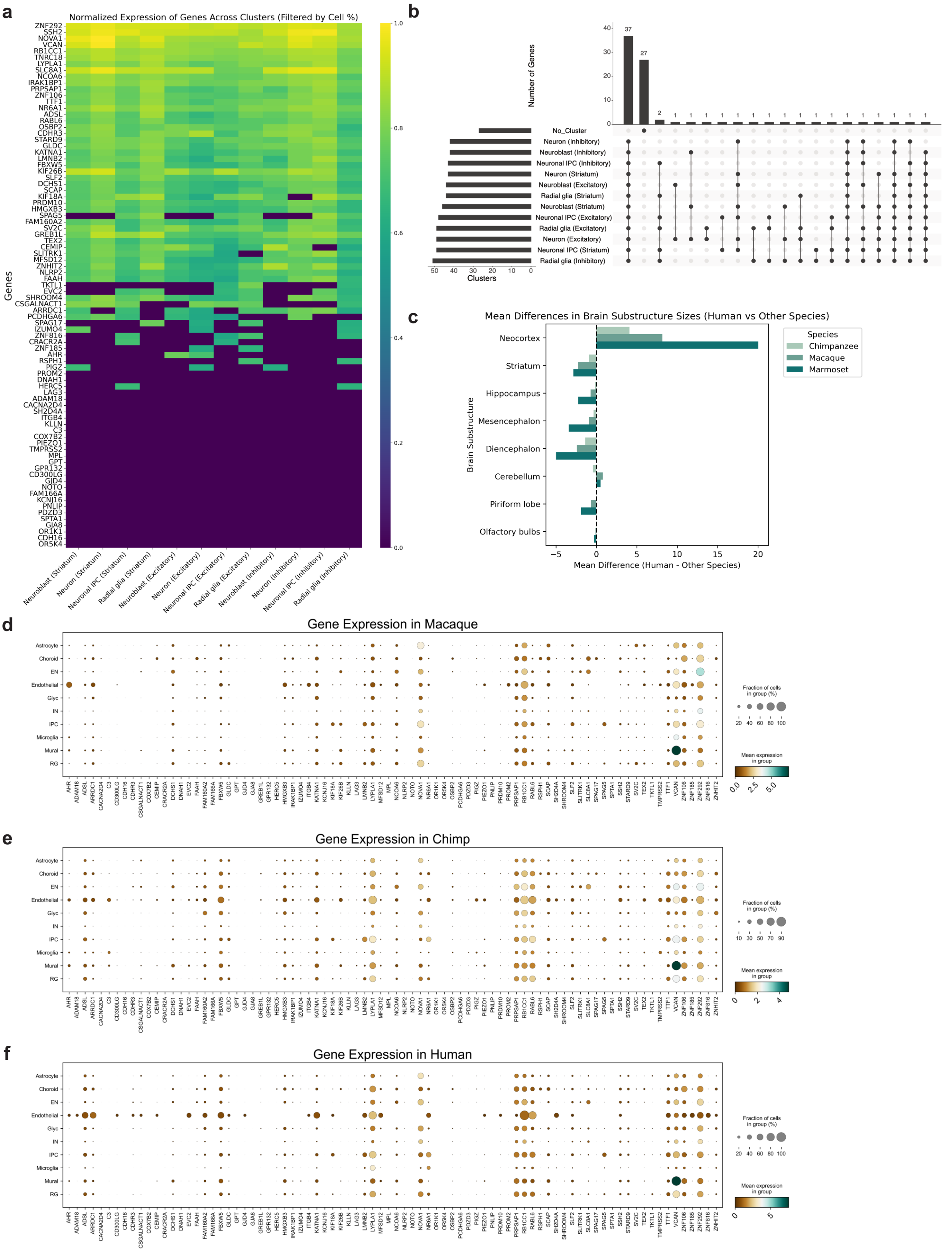


**Supplementary Figs. 1. Expression levels of non-synonymous modern human variants in the developing brain.** (a) Heatmap displaying the expression levels of 83 genes associated with non-synonymous variants in modern humans. Data were obtained from single-cell RNA sequencing (scRNA-seq) of first-trimester human brain cells, including radial glia, neuronal intermediate progenitor cells (IPCs), neuroblasts, and excitatory, inhibitory, and striatal neurons. (b) Upset plot illustrating the co-expression patterns of the 83 genes across different cell clusters, highlighting potential relationships between gene expression in specific cell populations. (c) Bar plot showing mean differences in brain structure sizes across primate species, represented as fold-change differences between humans and other primates. (d-f) Dot plots depicting the expression levels of the 83 genes associated with non-synonymous variants in macaque (d), chimpanzee (e), and human (f) scRNA-seq data. Dot color represents the average expression level, while dot size indicates the proportion of cells expressing each gene.

a

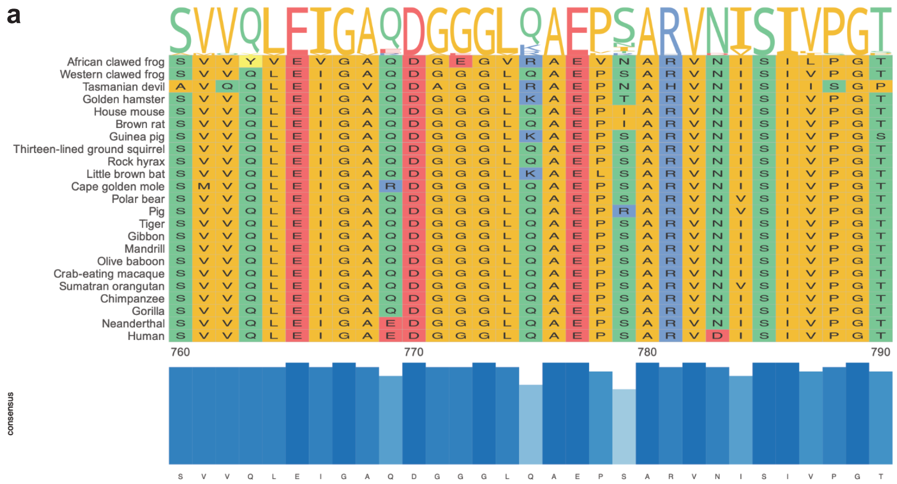


b

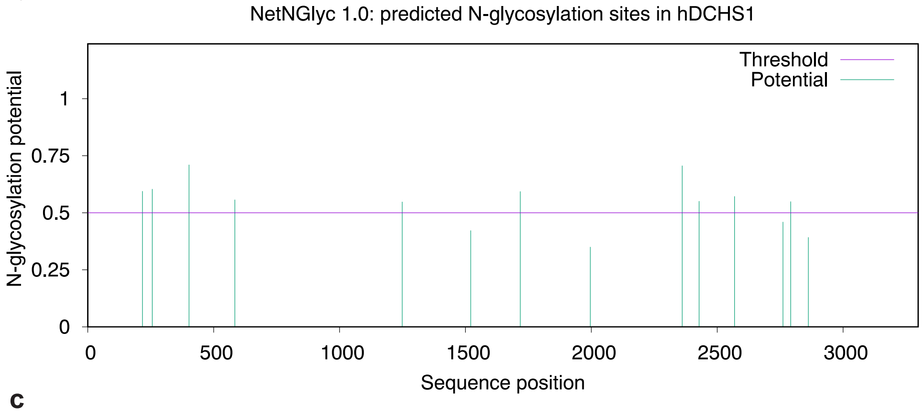


(Threshold=0.5)

| SeqName | Position | | Potential | Jury agreement | N-Glyc result |
|---|---|---|---|---|---|
| hDCHS1 | 217 | NRSH | 0.5949 | (6/9) | + |
| hDCHS1 | 256 | NQSR | 0.6041 | (8/9) | + |
| hDCHS1 | 402 | NVSL | 0.7107 | (9/9) | ++ |
| hDCHS1 | 584 | NASL | 0.5574 | (5/9) | + |
| hDCHS1 | 1249 | NGTI | 0.5482 | (7/9) | + |
| hDCHS1 | 1521 | NASR | 0.4224 | (8/9) | - |
| hDCHS1 | 1718 | NLTV | 0.5936 | (8/9) | + |
| hDCHS1 | 1996 | NASI | 0.3502 | (8/9) | - |
| hDCHS1 | 2361 | NLTV | 0.7066 | (9/9) | ++ |
| hDCHS1 | 2428 | NGTL | 0.5511 | (7/9) | + |
| hDCHS1 | 2569 | NLTV | 0.5721 | (8/9) | + |
| hDCHS1 | 2761 | NSST | 0.4601 | (5/9) | - |
| hDCHS1 | 2792 | NLSA | 0.5492 | (4/9) | + |
| hDCHS1 | 2862 | NQTT | 0.3926 | (7/9) | - |

c

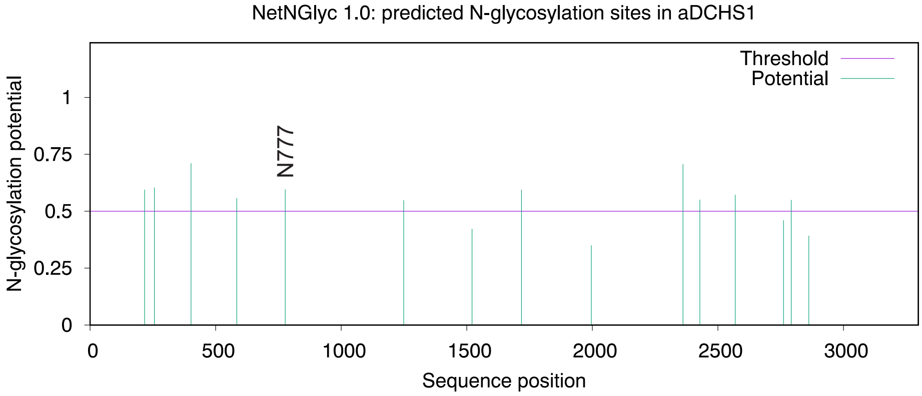


(Threshold=0.5)

| SeqName | Position | | Potential | Jury agreement | N-Glyc result |
|---|---|---|---|---|---|
| aDCHS1 | 217 | NRSH | 0.5949 | (6/9) | + |
| aDCHS1 | 256 | NQSR | 0.6041 | (8/9) | + |
| aDCHS1 | 402 | NVSL | 0.7107 | (9/9) | ++ |
| aDCHS1 | 584 | NASL | 0.5574 | (5/9) | + |
| aDCHS1 | 777 | NISI | 0.5959 | (8/9) | + |
| aDCHS1 | 1249 | NGTI | 0.5482 | (7/9) | + |
| aDCHS1 | 1521 | NASR | 0.4224 | (8/9) | - |
| aDCHS1 | 1718 | NLTV | 0.5936 | (8/9) | + |
| aDCHS1 | 1996 | NASI | 0.3501 | (8/9) | - |
| aDCHS1 | 2361 | NLTV | 0.7066 | (9/9) | ++ |
| aDCHS1 | 2428 | NGTL | 0.5511 | (7/9) | + |
| aDCHS1 | 2569 | NLTV | 0.5721 | (8/9) | + |
| aDCHS1 | 2761 | NSST | 0.4601 | (5/9) | - |
| aDCHS1 | 2792 | NLSA | 0.5492 | (4/9) | + |
| aDCHS1 | 2862 | NQTT | 0.3923 | (7/9) | - |

d

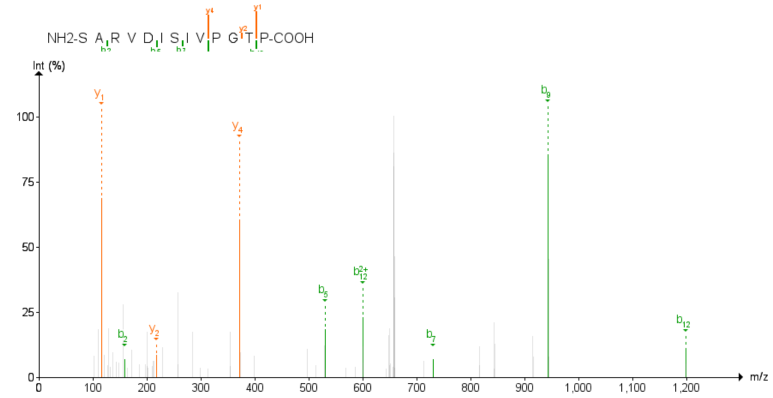


**Supplementary Figs. 2. Glycosylation at position 777 is absent in hDCHS1 and affects protein stability.** (a) Sequence alignment of the region encompassing the D777N mutation across eukaryotes. (b–c) *In silico* prediction of glycosylation sites in (b) human DCHS1 and (c) ancestral DCHS1 using the NetNGlyc 1.0 prediction score. Asparagine 777 in aDCHS1 is highlighted as a potential glycosylation site. (d) Mass spectrometry (MS) spectra of the hDCHS1 peptide corresponding to aspartic acid 777, confirming the absence of glycosylation as expected.

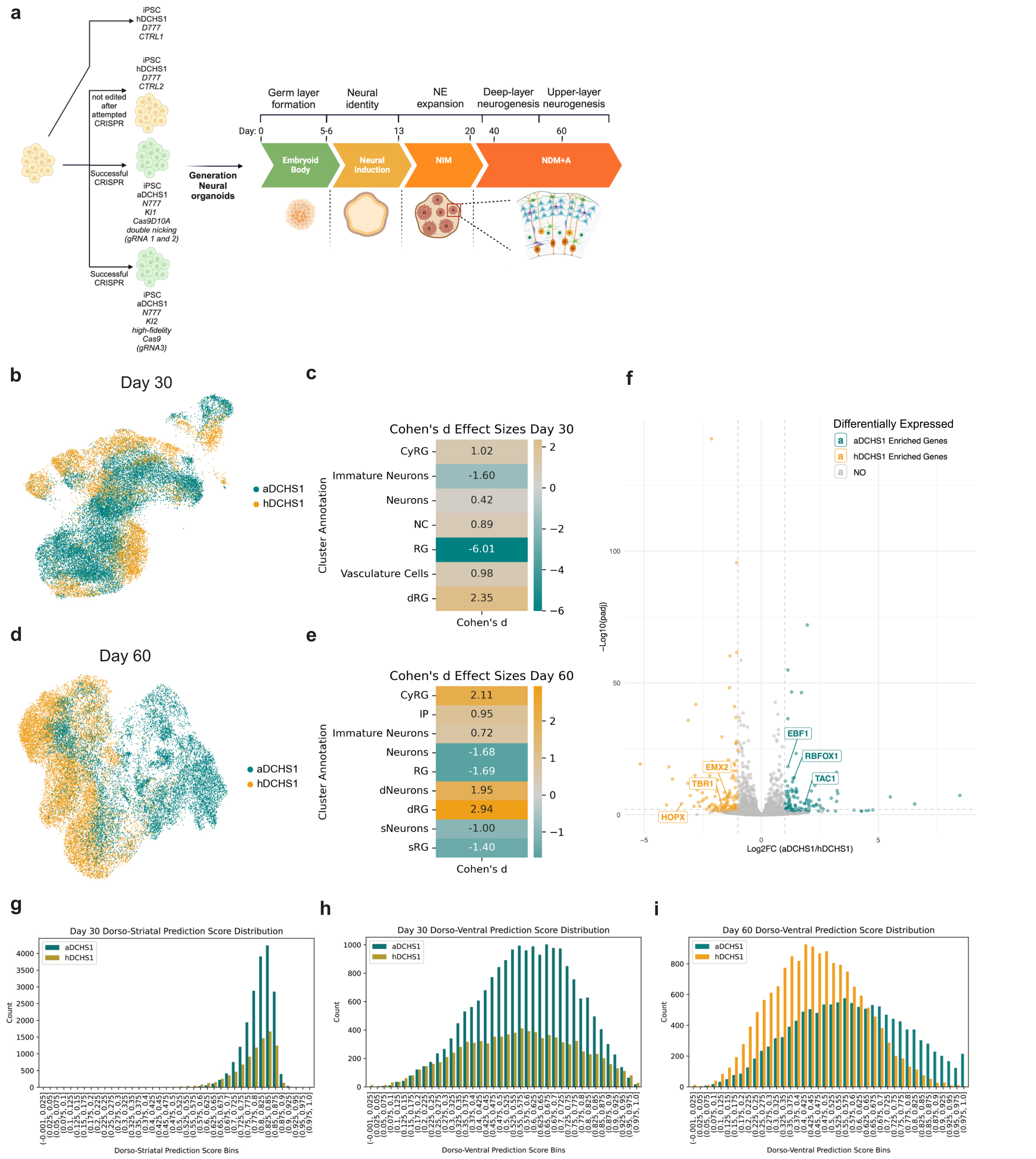


**Supplementary Figs. 3. CRISPR/Cas9 generation of aDCHS1 and hDCHS1 iPSCs reveals differences in neural organoid composition and dorsoventral patterning.** (a) Schematic representation of the rationale for generating aDCHS1 lines using CRISPR/Cas9. Two independent CRISPR-modified lines were generated for both aDCHS1 and hDCHS1. Created in https://BioRender.com. (b, d) Uniform manifold approximation and projection (UMAP) visualization of scRNA-seq data from hDCHS1 and aDCHS1 neural organoids at day 30 (b) and day 60 (d). (c, e) Cohen's *d* effect sizes for each cluster at day 30 (c) and day 60 (e), where positive values indicate a higher proportion in hDCHS1, while negative values indicate a skew toward aDCHS1. (f) Pseudobulk RNA-seq analysis of radial glia (RG) cells at day 30, comparing aDCHS1 and hDCHS1. (g) Histogram showing the dorso-striatal (DS) score of aDCHS1 and hDCHS1 at day 30, derived from first-trimester human embryonic telencephalon scRNA-seq data. (h) Histogram displaying the dorso-ventral (DV) score of aDCHS1 and hDCHS1 at day 30, based on neural organoid data. (i) Histogram showing the dorso-ventral (DV) score of aDCHS1 and hDCHS1 at day 60, based on neural organoid data.

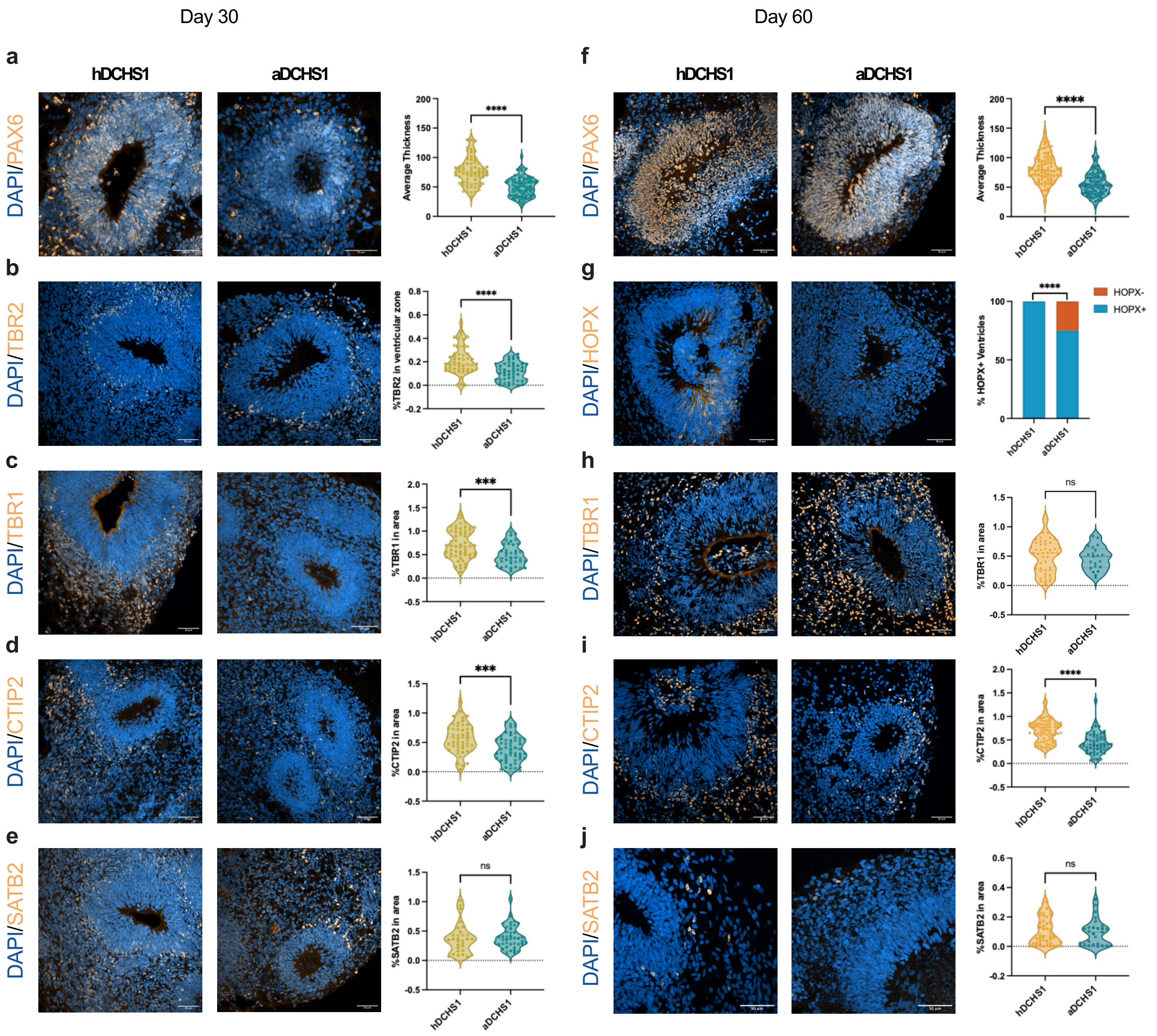


**Supplementary Figs. 4. Immunohistochemical characterization of cortical lineage markers in aDCHS1 and hDCHS1 neural organoids.** Representative immunohistochemistry (IHC) images of PAX6+ (a), TBR2+ (b), TBR1+ (c), CTIP2+ (d), and SATB2+ (e) cells in 30-day-old neural organoids, and PAX6+ (f), HOPX+ (g), TBR1+ (h), CTIP2+ (i), and SATB2+ (j) cells in 60-day-old hDCHS1 and aDCHS1 neural organoids. Relative quantification is shown as violin plots, with the median represented by a dotted line (*n* = number of analyzed ventricles from two independent batches of neural organoids, with four organoids analyzed per batch). Each shape represents an individual cell line. Statistical significance was assessed using one-way ANOVA. Scale bar: 50 μm.

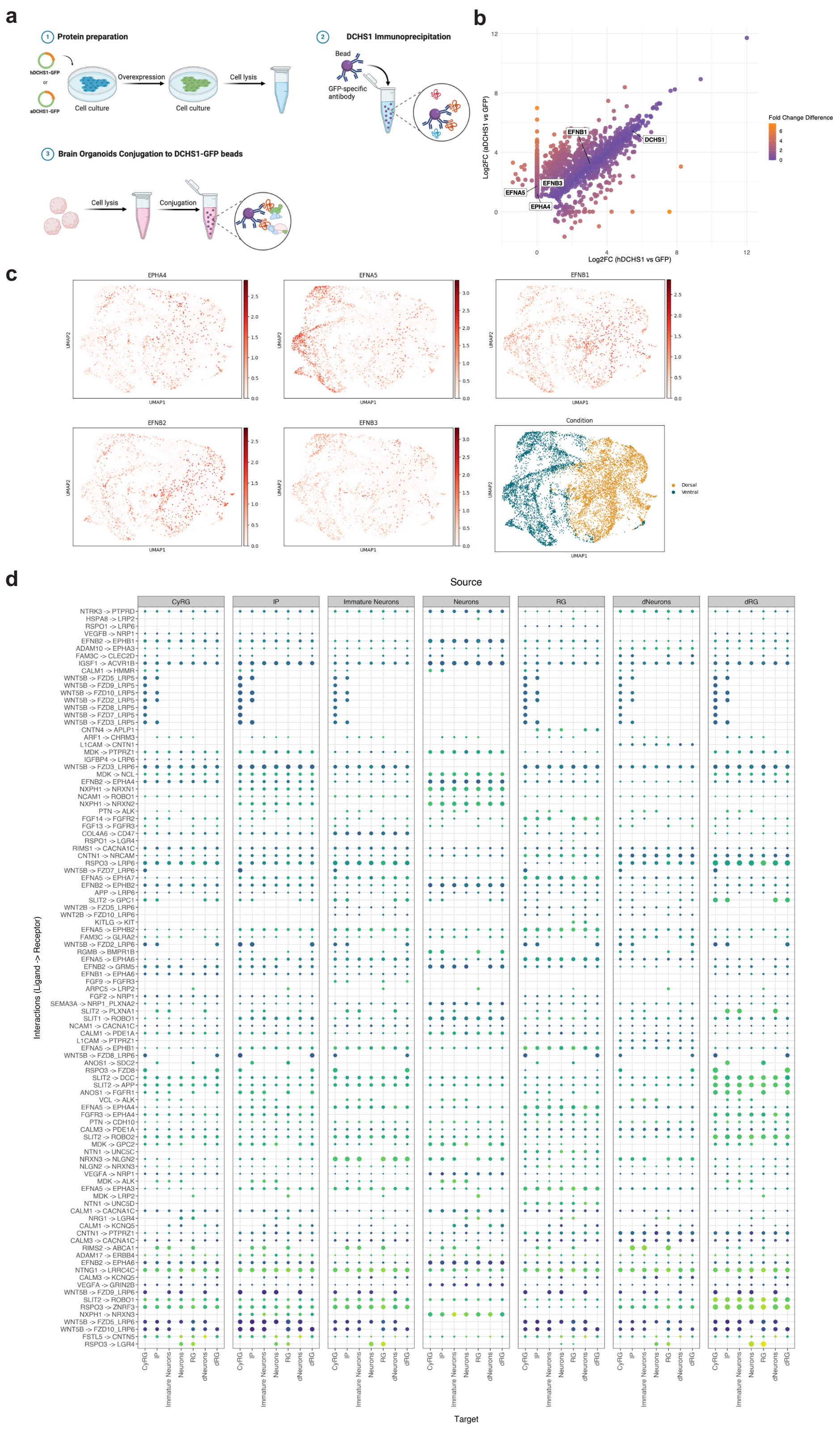


**Supplementary Figs. 5. Interactome analysis of ancestral and modern human DCHS1 reveals altered affinity for EPHA4 and its ligands.** (a) Schematic representation of the interactome experiment. TurboGFP-tagged hDCHS1 and aDCHS1 were overexpressed in HEK293 cells and immunoprecipitated using TurboGFP beads. The immunoprecipitated (IPed) DCHS1 was then incubated with lysates from 30-day-old neural organoids to assay for interacting proteins. DCHS1 interactors were identified via mass spectrometry. Created in https://BioRender.com. (b) Correlation plot of Log2 fold change (Log2FC) values comparing aDCHS1/GFP (y-axis) and hDCHS1/GFP (x-axis), showing only proteins significantly enriched in aDCHS1 and hDCHS1 immunoprecipitation experiments (Log2FC > 1, p-value < 0.05) and their relative expression. (c) UMAP projections displaying the expression patterns of *EPHA4*, *EFNA5*, *EFNB1*, and *EFNB3* in neural organoids.The rightmost panel shows the condition labels, distinguishing between dorsally and ventrally derived neural organoids. (d) Differential expression analysis of cell-cell communication events potentially deregulated between aDCHS1 and hDCHS1. The top 100 ligand-receptor interactions are shown. Scale bar: 50 µm.
