## Supplementary material for "DCHS1 Modulates Forebrain Proportions in Modern Humans via a Glycosylation Change": Tables

**Table 1 |** Primary antibodies

| **Antibody** | **Conc.** | **HIER performed** | **Source** | **Cat#** | **RRID** |
| --- | --- | --- | --- | --- | --- |
| Anti-CTIP2, rat monoclonal | 1:500 | yes, pH 6 or pH 9 | Abcam | ab18465 | AB_2064130 |
| Anti-FOXP2, mouse IgG1 monoclonal | 1:100 | no | Santa Cruz Biotechnology | sc-517261 | AB_2721204 |
| Anti-MEIS1/2/3, mouse IgG1 monoclonal | 1:300 | no | Santa Cruz Biotechnology | sc-101850 | AB_2143143 |
| Anti-PAX6, rabbit polyclonal | 1:250 | yes, pH 6 | BioLegend | 901302 | AB_2565003 |
| Anti-EOMES, rabbit polyclonal | 1:500 | yes, pH 6 | abcam | ab23345 | AB_778267 |
| Anti-DCHS1, rabbit polyclonal | 1:100 | no | Thermo Fisher Scientific | PA5-85905 | AB_2802706 |
| Anti-EPHA4, mouse IgG1 | 1:100 | no | Thermo Fisher Scientific | 37-1600 | AB_2533301 |
| Anti-TBR1, rabbit polyclonal | 1:500 | no | Abcam | ab31940 | AB_2200219 |
| Anti-SATB2, monoclonal | 1:500 | yes, pH 6 | Abcam | ab51502 | AB_882455 |
| Anti-HOPX, rabbit polyclonal | 1:1000 | yes, pH 6 | Atlas Antibodies | HPA030180 | AB_10603770 |

**Table 2 |** Secondary antibodies

| **Antibody** | **Conc.** | **Source** | **Cat#** | **RRID** |
| --- | --- | --- | --- | --- |
| Alpaca anti-Mouse IgG1 Nanobody, Alexa Fluor 488 | 1:500 | ChromoTek | sms1AF488-1 | AB_2827578 |
| Alpaca anti-Mouse IgG1 Nanobody, Alexa Fluor 568 | 1:500 | ChromoTek | sms1AF568-1 | AB_2827578 |
| Goat anti-Mouse IgG (H+L), Alexa Fluor 488 | 1:500 | Thermo Fisher Scientific | A-11001 | AB_2534069 |
| Goat anti-Mouse IgG (H+L), Alexa Fluor 546 | 1:500 | Thermo Fisher Scientific | A-11003 | AB_2534071 |
| Goat anti-Rabbit IgG (H+L), Alexa Fluor 546 | 1:500 | Thermo Fisher Scientific | A-11010 | AB_2534077 |
| Goat anti-Rabbit IgG (H+L), Alexa Fluor 647 | 1:500 | Thermo Fisher Scientific | A-21244 | AB_2535812 |
| Goat anti-Rat IgG (H+L), Alexa Fluor 647 | 1:500 | Thermo Fisher Scientific | A-21247 | AB_141778 |
